# Cryptochrome 2 is necessary for promoting a blue light-independent chloroplast signal to control cellular degradation

**DOI:** 10.64898/2025.12.01.690775

**Authors:** David W. Tano, Kyle Palos, Karen E. Fisher, Robert A. Easter, Andrew D. L. Nelson, Jesse D. Woodson

**Affiliations:** The School of Plant Sciences, University of Arizona, Tucson, AZ; Boyce Thompson Institute, Ithaca, NY 14853

## Abstract

To thrive in dynamic environments, plants can use their photosynthetic plastid (chloroplast) organelles to sense environmental changes and relay this information to the cell. This signaling can be initiated within chloroplasts by the accumulation of reactive oxygen species (ROS) such as singlet oxygen (^1^O_2_), which can accumulate under stress and initiate retrograde (chloroplast-to-nucleus) signaling, chloroplast degradation, and programmed cell death (PCD). However, the mechanisms driving these signals, and the relationship between gene expression and cellular degradation, are unknown. To understand this physiology, here we use the *Arabidopsis thaliana plastid ferrochelatase 2* (*fc2*) mutant, which conditionally produces chloroplast ^1^O_2_ and initiates signaling. We show that the blue light photoreceptor cryptochrome 2 (CRY2) is necessary to promote ^1^O_2_-induced chloroplast degradation and PCD in the *fc2* mutant and that *cry2* blocks cellular degradation without affecting bulk ^1^O_2_ levels, suggesting that signaling, rather than photo-oxidative stress, has been blocked. Furthermore, a global RNA transcript analysis demonstrated that the impact of the *cry2* mutation on retrograde signaling is limited and that the effect of ^1^O_2_ on nuclear gene expression and cellular degradation can be uncoupled. Under permissive conditions, *fc2 cry2* mutants exhibited constitutive expression of photo-oxidative stress genes, suggesting that *cry2* may lead to transcriptional priming to photo-oxidative stress. Finally, we demonstrate that this function of CRY2 is independent of blue light-signaling and that CRY2 isoforms lacking canonical blue light signaling capabilities restore ^1^O_2_-induced PCD to *fc2 cry2*. Together, these results reveal a blue light-independent role for CRY2 in chloroplast signaling and stress acclimation pathways.

## Introduction

As sessile organisms rooted to the ground, plants must utilize molecular mechanisms that allow them to swiftly respond to stresses and changes in their environments. Energy producing organelles (i.e., photosynthetic plastids [chloroplasts] and mitochondria) play important roles in this ability and can serve as environmental sensors for plant cells. As sites of high-energy redox chemistry for photosynthesis and respiration, these organelles are particularly sensitive to environmental changes and stresses, which can lead to rapid dysfunction and the subsequent production of reactive oxygen species (ROS) and other damaging metabolites [1, 2]. This is particularly true for chloroplasts in light-exposed photosynthetic tissues, which can quickly produce large amounts of singlet oxygen (^1^O_2_) and superoxide (O_2_^-^) (which can be subsequently converted to hydrogen peroxide [H_2_O_2_] by superoxide dismutase) at photosystems II and I, respectively [3], under multiple stresses including excess (high) light [4], drought [5], salinity [6], heat [7],and pathogen attack [8]. Chloroplasts contain a suite of ROS detoxifying enzymes (e.g., peroxidases) and molecules (e.g., tocopherols), but these defenses can become overwhelmed under severe stress, allowing ROS to rapidly oxidize nearby macromolecules such as DNA, RNA, proteins and lipids [3]. At the same time, these ROS are sources of information about a changing environment and act as signaling molecules to reprogram hundreds of nuclear genes (via retrograde [chloroplast-to-nucleus] signals) to mitigate stress, improve photosynthetic efficiency, or coordinate chloroplast development and/or turnover [9–13].

In addition to retrograde signaling, ^1^O_2_ induces cellular degradation in the form of chloroplast degradation (aka, chloroplast quality control) and programmed cell death (PCD) [12]. It is hypothesized that the degradation of dysfunctional chloroplasts ensures a healthy population of chloroplasts, initiating PCD limits further necrosis and water loss, and both processes allow cells to recycle the nitrogen-rich content of chloroplasts [13–15]. ^1^O_2_ has a very short half-life (∼0.5–1.0 μs), and its limited diffusion radius (∼200 nm) makes it unlikely to leave the relatively large chloroplasts (∼2-10 μm wide) in which it is made [16]. As such, any ^1^O_2_ signal to control retrograde signaling and cellular degradation must rely on secondary messengers to relay information from the chloroplast to the nucleus and/or cell. To identify such factors, researchers have used *Arabidopsis thaliana* mutants that conditionally accumulate ^1^O_2_ and activate such signaling. One mutant, *fluorescent in blue light* (*flu*), over-accumulates the chlorophyll precursor protochlorophyllide (Pchlide) in the dark [17]. Upon light exposure, Pchlide is photo-excited and promotes the production of ^1^O_2_. This rapid accumulation of ^1^O_2_ induces retrograde signaling and is followed by PCD [18]. A forward genetic screen identified the chloroplast-localized protein Executor 1 (EX1) as necessary for these responses, demonstrating for the first time that ^1^O_2_-induced PCD is a genetically controlled response to chloroplast dysfunction [19].

Another mutant, *plastid ferrochelatase 2* (*fc2*), also conditionally accumulates ^1^O_2_, albeit by a different mechanism [20, 21]. When *fc2* is grown under diurnal light cycling conditions, the heme/chlorophyll precursor and FC2 substrate protoporphyrin IX (Proto) accumulates within minutes after dawn. Photo-excitation of Proto then leads to the production of ^1^O_2_, which signals rapidly to induce the expression of hundreds of genes (∼1 hour), followed by chloroplast degradation (∼2-3 hours), and eventually PCD (∼8 hours) [20–22]. When grown under permissive constant light conditions, *fc2* plants do not experience PCD. However, low levels of Proto and ^1^O_2_ do accumulate and promote selective chloroplast degradation, typified by chloroplasts being internally degraded in the cytoplasm before interacting with the central vacuole in a process resembling fission-type microautophagy [23, 24]. To identify factors involved in ^1^O_2_ signaling in *fc2*, a forward genetic screen identified mutants with blocked ^1^O_2_-induced PCD and chloroplast degradation, which were dubbed *ferrochelatase two suppressor* (*fts*) mutants. The identification of the affected genes revealed a signaling role for plastid gene expression [22, 25] and chloroplast ubiquitination [20] and led to a model where a plastid-encoded gene product(s) is required to transmit a ^1^O_2_ signal from chloroplasts, which leads to the recruitment of ubiquitination machinery to mark chloroplasts for turnover [26]. These mutations also blocked retrograde signaling to the nucleus, suggesting that cellular degradation and nuclear gene expression originate from the same signal.

Unexpectantly, the ^1^O_2_ signals induced in *fc2* and *flu* appear to represent different pathways. A comparative study and meta-analysis of ^1^O_2_-induced gene expression revealed that a similar set of nuclear genes were controlled by ^1^O_2_ in these mutants (39% of genes induced in *fc2* were also induced in *flu*) despite the plants being grown under very different conditions [27]. However, a genetic analysis suggested that the pathways induced in these two backgrounds utilize different mechanisms [20, 27]. For example, *ex1*, which blocks ^1^O_2_-induced PCD in *flu*, was unable to block ^1^O_2_-induced PCD in *fc2*. Conversely, *pub4*, which blocks ^1^O_2_-induced PCD in *fc2*, was unable to block ^1^O_2_-induced PCD in *flu*. Together, these results suggest that multiple ^1^O_2_ signaling pathways exist to control cellular degradation. The factors involved in these signals, particularly those acting outside the chloroplast, are poorly understood and represent a major knowledge gap in plant cell biology.

Plants also sense their environments using a suite of photoreceptors which are activated under different wavelengths of light to control nuclear gene expression, photomorphogenesis, chloroplast function, and many other aspects of plant development [28]. Two groups of photoreceptors, phytochromes and cryptochromes, have been shown to impact multiple types of chloroplast signals, suggesting that they could also be important for mediating chloroplast-induced cellular degradation pathways. Phytochrome A (phyA) and phytochrome B (phyB) perceive far-red (660 nm) and red light (730 nm), respectively, and have been shown to affect chloroplast signals to coordinate the expression of photosynthesis associated nuclear genes (PhANGs) with the developmental state of the chloroplast (phyA and phyB) [29] and to be involved in transmitting methylerythritol cyclodiphosphate (MEcPP)-dependent chloroplast stress signals (phyB) [30], which can be activated under high light and drought stress.

In plants, two main cryptochromes (cryptochrome 1 [CRY1] and CRY2) sense blue light (320-500 nm) to control photomorphogenesis, chloroplast development, and flowering initiation [31]. Photo-activation of CRYs depends on a flavin adenine dinucleotide (FAD) chromophore, which is reduced by blue light irradiance. This causes CRYs to undergo conformational changes and homeroligomerization, which is necessary for CRYs to perform their canonical light signaling functions via protein interactions to regulate gene expression and development in the nucleus (and in the case of CRY1, also the cytoplasm) [32, 33]. Both CRYs play overlapping functions with respect to photomorphogenesis, albeit at different fluence rates of blue light: CRY1 plays a dominant role under high fluence rates, while the function of CRY2 is limited to lower fluence rates (< 10 µmol photons m^-2^ sec^-1^). Importantly, CRY1 has been implicated in several chloroplast retrograde signaling pathways and has been shown to affect the coordination of PhANGs in response to chloroplast development [29], to propagate signals from chloroplasts under high light stress [34], and as part of the chloroplast ^1^O_2_ signal to control PCD in the *flu* mutant [35]. In the latter case, the ^1^O_2_ signal in *flu* protoplasts was shown to be blue light-dependent and blocked in *cry1* mutants. As CRY1 is localized to the cytoplasm and nucleus, it was suggested that CRY1 plays a signaling role downstream of chloroplast-localized EX1. CRY2 played a minimal role in controlling this PCD, and to the best of our knowledge, has not been implicated in any chloroplast stress signaling.

Here, we tested the roles of phytochromes and CRYs in ^1^O_2_ signaling in *fc2* and demonstrate that CRY2 plays a robust role in promoting ^1^O_2_ controlled PCD and chloroplast degradation. Its impact on retrograde signaling, however, is limited and *fc2 cry2* mutants still misregulate most genes affected by ^1^O_2_. Instead, the *cry2* mutation leads to the constitutive induction of a subset of stress-related genes, which may allow *fc2 cry2* plants to acclimate to photo-oxidative stress. Finally, we demonstrate that this function of CRY2 is independent of blue light signaling and that isoforms incapable of being photo-activated or relaying canonical blue light signaling can complement ^1^O_2_-induced PCD in *fc2 cry2* mutants. Together, these results point to a blue light-independent role for CRY2 in chloroplast signaling to control a plant’s response to photo-oxidative damage.

## Results

### The photoreceptor CRY2 is necessary for singlet oxygen-induced cellular degradation in *fc2* mutants

It was previously reported that the blue light photoreceptor CRY1 is required to transmit the chloroplast ^1^O_2_ stress signal for PCD in *flu* mutants [35]. We wanted to determine if photoreceptors also play a ^1^O_2_-signaling role in the *fc2* mutant. To this end, we crossed *cry1*, *cry2-1* (hereafter referred to as *cry2*), *phyA*, and *phyB* mutations into the *fc2* background to test their ability to suppress ^1^O_2_-induced PCD under cycling light conditions. Mutant combinations were confirmed via genotyping and by measuring hypocotyl elongation under monochromatic LED light sources. As expected, *phyA*, *phyB*, and *cry1*/*cry2* mutations led to increased hypocotyl lengths (compared to wt and *fc2*) in far-red, red, and blue light, respectively (**Figs. S1a-c**). The *fc2* background had no significant effect on these phenotypes suggesting that light signaling occurs normally in this mutant. Next, we tested the ability of these mutations to block ^1^O_2_-induced PCD. When seedlings were grown under 24h (constant) light conditions, all genotypes appeared healthy (**Figs. 1a** and **S1d**), although lines with the *fc2* mutation appeared slightly pale and accumulated less total chlorophyll (**Figs. 1b and S1e**). When seedlings were grown under diurnal 6h white light / 18h dark (cycling) conditions, *fc2* seedlings bleached, accumulated less total chlorophyll, and died in a manner resembling PCD (**Figs. 1a and b and S1d and e**). This response was reversed in *fc2 pub4*, confirming that the *fc2* PCD phenotype was caused by ^1^O_2_-signaling (**Fig. S1d**) [20]. Neither *phyA* or *phyB* reversed the PCD phenotype and *fc2 phyA* and *fc2 phyB* were indistinguishable from *fc2* (**Figs. S1d and e**). As previously reported [27], *cry1* also did not reverse the *fc2* PCD phenotype (**Fig. 1a**). However, *fc2 cry2* mutants appeared healthy under cycling light conditions, suggesting that *cry2* was blocking ^1^O_2_-induced PCD (**Fig. 1a**). These phenotypes were confirmed with a trypan blue stain to assess cell death. As expected from the visual phenotypes, only *cry2* and *pub4* significantly decreased cell death in *fc2* under cycling light conditions (**Figs. 1c and d and S1f and g**). We also tested the phenotype of a *fc2 cry1 cry2* triple mutant. This line also survived under cycling light conditions, but the level of PCD was comparable to the *fc2 cry2* mutant, suggesting that the *cry1* mutation did not have an additive effect on stress tolerance (**Figs. 1a, c, and d**). As such, we only moved forward with the *fc2* photoreceptor double mutants.

**Figure 1.**
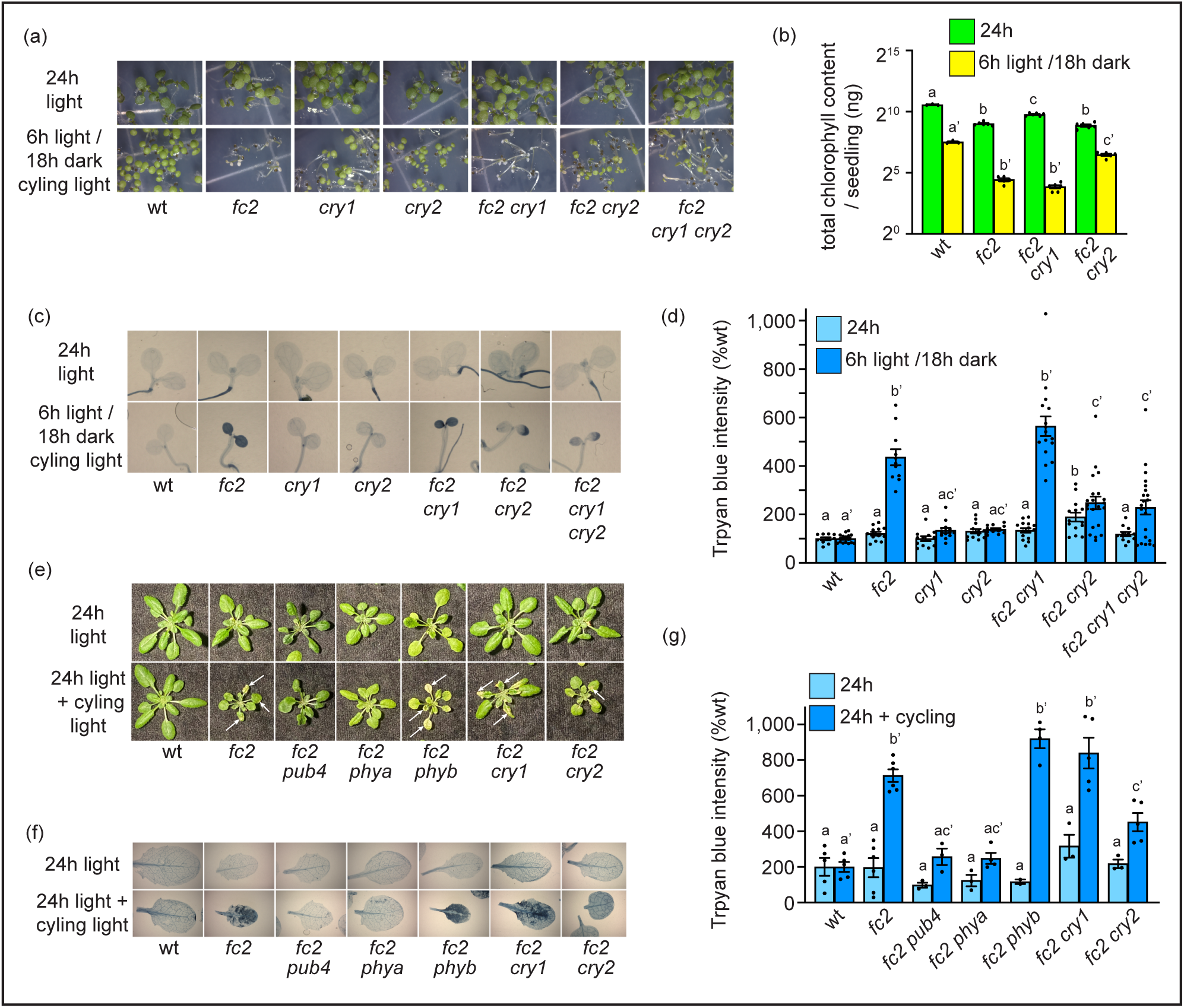
The blue light photoreceptor CRY2 s necessary for singlet oxygen-induced cell death in *fc2*. The effect of the *cry2* mutation on singlet oxygen-induced programmed cell death in the *fc2* mutant was tested. **A)** Shown are six-day-old seedlings grown under constant white light (24h) or diurnal cycling white light (6h white light / 18h dark) conditions. **B)** Mean levels of total chlorophyll (per seedling) of seven-day-old seedlings grown in constant white light (24h) (+/- SE, n = 6 replicates). **C)** Shown are representative trypan blue stains of seedlings in panel A. The dark blue color is indicative of cell death. **D)** Shown are mean intensities of trypan blue (+/- SE, n ≥ 4 seedlings) from **C**. **E)** Shown are representative 23-day-old plants grown under constant white light (24h) or under 24h white light for 17 days and then shifted to diurnal cycling white light conditions (16h white light / 8h dark) for 6 additional days. Arrows indicate lesions. **F)** Shown are representative trypan blue cell death stains of leaves from plants in panel E. The dark blue color is indicative of cell death. **G)** Shown are mean intensities of trypan blue (+/- SE, n ≥ 3 leaves from different plants) from **F**. For panels B, D, and G, statistical analyses were performed by a one-way ANOVA test followed by Tukey’s HSD. Letters indicate statistically significant differences between samples (*P* ≤ 0.05). Separate analyses were performed for the different light conditions, and the significance of the cycling light is denoted by letters with a prime symbol (ʹ). Closed circles represent individual data points.

The ability to suppress ^1^O_2_-induced PCD can be life stage specific [27, 36], leading us to test phenotypes in adult plants. When grown under constant white light conditions for 23 days, all lines appeared healthy (**Fig. 1e**). However, if 17-day-old plants were shifted to cycling light conditions (16h white light / 8h dark) for six days, the *fc2* mutant accumulated visible lesions on its leaves (**Fig. 1e**), which was confirmed to be PCD with a trypan blue stain (**Figs. 1f and g**). PCD was mostly absent in *fc2 pub4* mutants, which confirmed that that the lesions were ^1^O_2_ signaling dependent. As in seedlings, *cry2* also reduced the level of PCD compared to *fc2*, albeit to a lesser degree. Surprisingly, *fc2 phyA* mutants had less PCD than *fc2*, suggesting that *phyA* plays a stage-specific role in promoting ^1^O_2_-signaling. Both *cry1* and *phyB* failed to suppress PCD, confirming neither protein plays a prominent role in ^1^O_2_-signaling in *fc2*, regardless of life stage. Together these results suggest that phyA may play a specialized ^1^O_2_-signaling role in adult tissue to control cellular degradation, but CRY2 plays a more robust role, regardless of life stage. As such, we decided to continue our focus on the *cry2* mutation.

To confirm that the *cry2* mutation was causative for the suppression of PCD in *fc2* mutants, we took three approaches. First, we performed a genetic linkage experiment by crossing *cry2* with *fc2* and testing if the *cry2* lesion was linked to suppression of PCD in *fc2*. F2 plants were genotyped for the *fc2* mutation and 43 *fc2*/*fc2* plants (with *cry2* independently segregating) were transplanted to soil. The *cry2* late flowering phenotype [37] was then scored in these plants and six with a clear late flowering phenotype (≥50 days post germination) were identified (**Fig. S2a**). F3 plants from all 43 F2 plants were then grown under cycling light conditions and rates of survival were scored. The six lines with late flowering phenotypes survived light cycling conditions, while the other 37 lines did not or had segregating phenotypes (**Fig. S2b**). Together, this result indicates that the *cry2* late flowering phenotype is closely linked to the suppression of PCD (X^2^ value = 2.80, critical value (p ≤ 0.05) = 3.84). Next, we tested if other *cry2* alleles can block ^1^O_2_-induced PCD in *fc2*. Three additional mutations were identified in the Landsberg erecta (Ler) background (*fha-1* [W54stop], *fha-2* [G254R], and *fha-3* [frameshift after R478]) and were crossed into a Ler *fc2* to create three Ler *fc2 cry2* mutants (**Fig. S3A**). These three alleles also reversed the *fc2* PCD phenotype under cycling light conditions confirming that they phenocopy the *cry2-1* mutation (**Figs. S3b-d**). Finally, we complemented *fc2 cry2* with a wt copy of *CRY2* driven by the constitutive *35S* promoter (*35S*::*CRY2*). Three independent lines exhibited restored PCD under light cycling conditions (**Fig. S4a-c**). Together, these results confirm the causality of the *cry2* mutation in suppressing PCD in *fc2*.

### CRY2 is required for chloroplast quality control pathways, but not for general chloroplast development

In the *fc2* mutant, rates of chloroplast degradation are increased in permissive constant light conditions even in the absence of PCD [24]. To test the effect of *cry2* on such cellular degradation, we used transmission electron microscopy (TEM) to assess chloroplast ultrastructure in four-day-old seedlings grown under constant light conditions. Most chloroplasts in *fc2* appeared structurally normal and resembled wt chloroplasts (**Fig. 2a**). However, approximately 40% of *fc2* chloroplasts appeared to be damaged and in a state of degradation, with disordered and compressed thylakoid membranes and large internal vesicles (**Fig. 2b**). This rate of degradation was significantly reduced by the *cry2* mutation and only 11% of chloroplasts in *fc2 cry2* exhibited some level of degradation (**Fig. 2c**). No degradation was observed in wt or *cry2* seedlings, and their chloroplasts appeared similar. Together, these results suggest that the *cry2* mutation specifically blocks chloroplast quality control pathways that direct chloroplast degradation in the absence of PCD.

**Figure 2.**
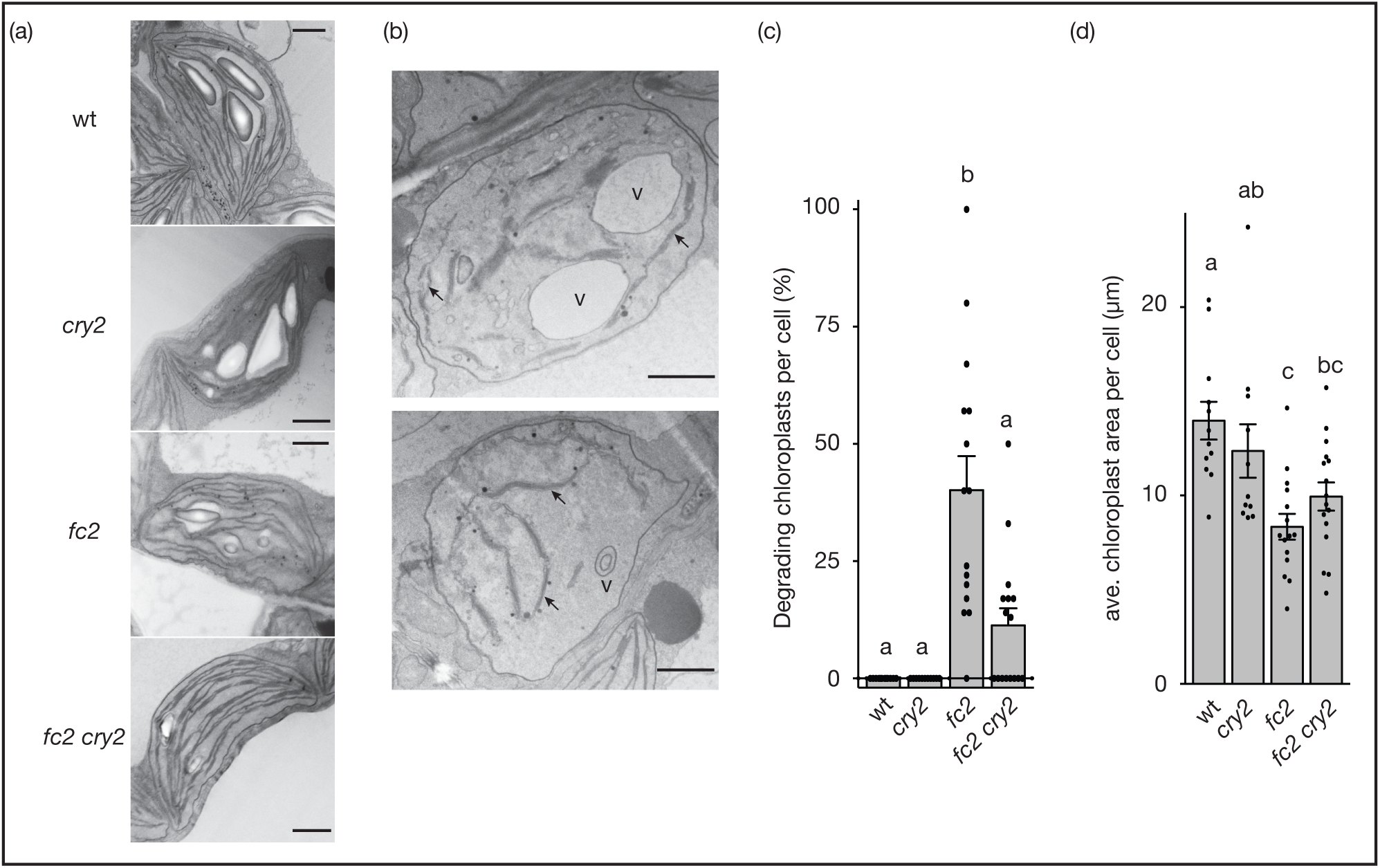
CRY2 is necessary for chloroplast quality control pathways in the *fc2* mutant. The effect of the *cry2* mutation on chloroplast degradation in the *fc2* mutant was tested. **A)** Shown are representative transmission electron microscopy images of chloroplasts in four-day-old seedlings grown under constant light conditions. **B)** Representative images of degrading chloroplasts in *fc2* in same seedlings used in A. Scale bars = 1 um. **C)** Shown are the average % of chloroplasts being degraded within a cell (+/- SE, n ≥ 11 cells). **D)** Average chloroplast area (um^2^) per cell (n ≥ 11 cells). For panels B and C, at least 3 seedlings were imaged. Statistical analyses were performed by a one-way ANOVA test followed by Tukey’s HSD. Letters indicate statistically significant differences between samples (*P* ≤ 0.05). Closed circles represent individual data points.

Several of the *fts* mutations that block ^1^O_2_-induced chloroplast degradation also reduce plastid gene expression and delay chloroplast development [20, 22, 25]. To test if *cry2* was blocking PCD and/or chloroplast degradation in a similar way, we first assessed chloroplast development in this mutant and measured the average area of chloroplasts per cell using the TEM images of chloroplasts in seedlings. As previously reported [24], *fc2* chloroplasts were slightly smaller than wt chloroplasts (**Fig. 2d**). However, *cry2* did not affect chloroplast size and *cry2* and *fc2 cry2* mutants had similar sized chloroplasts to wt and *fc2*, respectively. Next, we measured the ability of these mutants to accumulate Pchlide in the dark, which is an indicator of tetrapyrrole (e.g., chlorophyll) synthesis and chloroplast development [38]. As previously reported, *fc2* mutants accumulate 2-3-fold more Pchlide than wt [20, 39] (**Fig. S5a**). However, neither *cry1* or *cry2* reduced the accumulation of Pchlide in the wt or *fc2* backgrounds. We also measured the expression of six genes involved in tetrapyrrole synthesis: plastid *trnE* (the pathway precursor) and five nuclear genes encoding key regulatory steps (*HEMA1*, *HEMA2*, *CHLH*, *GUN4*, and *PORA*) [38]. None of the six genes were significantly repressed in *fc2 cry1* or *fc2 cry2* compared to *fc2* in constant light (**Fig. S5b**) or cycling light conditions (**Fig. S5c**). Instead, *trnEY* transcripts were increased in *fc2 cry2* in cycling light conditions and *PORA* transcripts were increased in both conditions. Finally, we tested if *cry2* affects plastid gene expression, which has been hypothesized to be a possible mechanism by which to reduce ^1^O_2_ signaling [22, 25]. We tested the expression of seven (*psbA*, *psbB*, *psbD*, *psaJ*, and *rbcL*) and two (*rpoB* and *clpP*) genes transcribed by the plastid-encoded RNA polymerase (PEP) and nuclear-encoded RNA polymerase (NEP), respectively. *cry1* and *cry2* did not significantly reduce the expression of any of these genes compared to *fc2* in constant light (**Fig. S5d**) or cycling light conditions (**Fig. S5e**). We also assessed expression of three *SIGMA FACTOR* genes necessary for plastid transcription [40], early chloroplast development (*SIG2* and *SIG6*) [41, 42], and blue light responses (*SIG5*) [43]. None of the *SIG* genes were repressed in *fc2 cry2* compared to *fc2* in constant light (**Fig. S5f**) or cycling light conditions (**Fig. S5g**). Conversely, all three genes were significantly repressed in *fc2 cry1* compared to *fc2* under cycling light conditions. Together, these results demonstrate that *cry2* does not block chloroplast ^1^O_2_ signaling by measurably delaying chloroplast development or reducing chloroplast gene expression.

### The *cry2* mutation uncouples programmed cell death from canonical retrograde signaling

Next, we aimed to determine what effects the *cry2* mutation has on ^1^O_2_ accumulation and its induction of canonical retrograde signaling. First, we measured bulk levels of ^1^O_2_ accumulation in seedlings grown under cycling light conditions using Singlet Oxygen Sensor Green (SOSG). As expected, ^1^O_2_ accumulated in *fc2* two hours after subjective dawn (**Figs. 3a and b**). Neither *cry1*, *cry2*, or the *cry1 cry2* combination reduced this ^1^O_2_ accumulation. Instead, the *cry1* mutation slightly, but significantly, increased ^1^O_2_ levels. Together, this suggests that *cry2* does not block ^1^O_2_ signaling by reducing bulk levels of this ROS.

**Figure 3.**
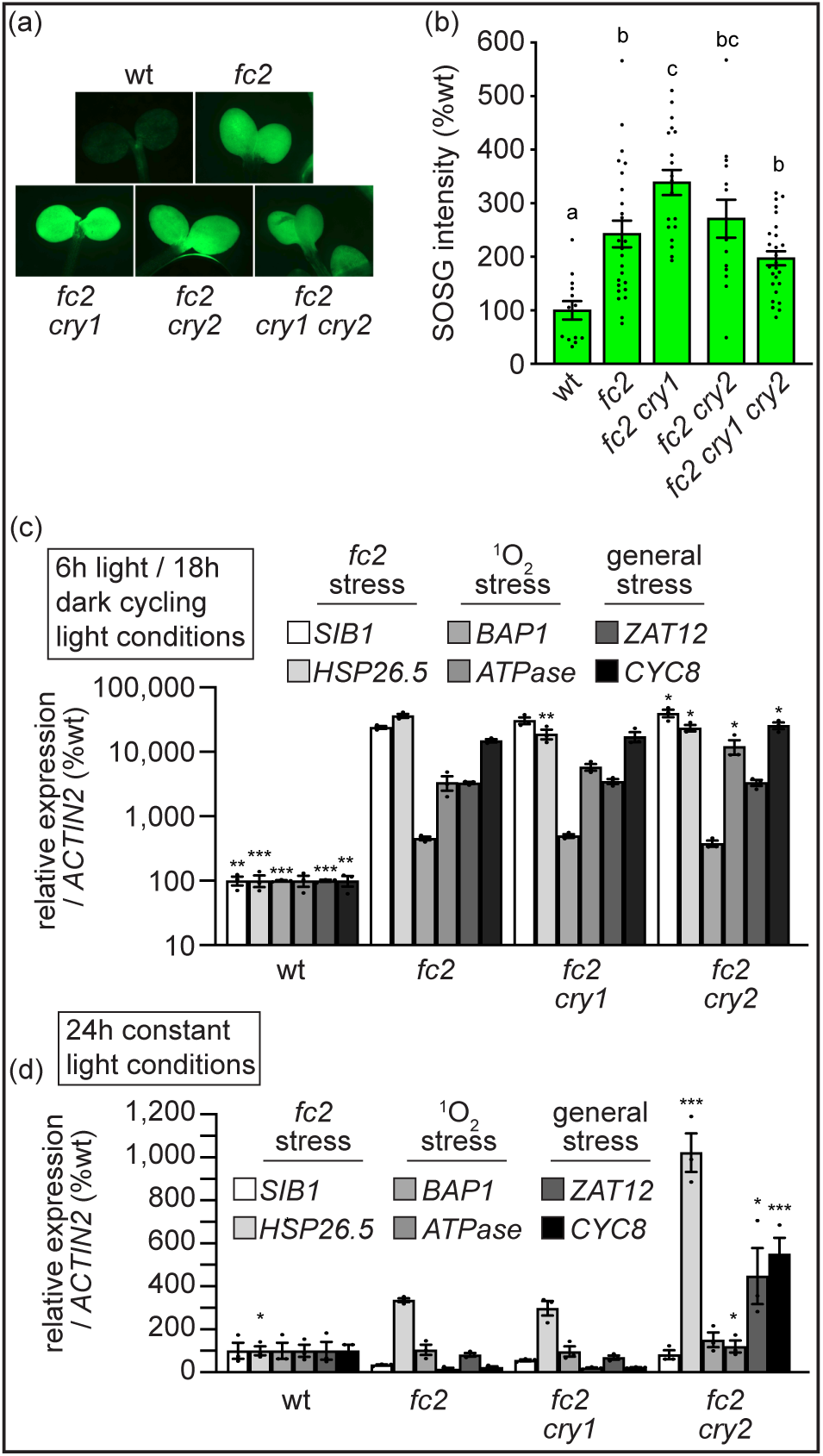
CRY2 function is independent of singlet oxygen accumulation and retrograde signaling in *fc2*. The *cry1* and *cry2* mutations were tested for their ability to affect singlet oxygen (^1^O_2_) accumulation and retrograde signaling. **A)** Shown are representative images of four-day-old seedlings stained with Singlet Oxygen Sensor Green (SOSG). Seedlings were grown for three days in 6h white light / 18h dark diurnal light cycling conditions, dark incubated at the end of day three, and re-exposed to light on day four. Images were acquired 3h post-dawn. **B)** Shown are mean SOSG intensities (+/- SE, n ≥ 13 seedlings) from panel A. RT-qPCR analysis of stress marker gene transcripts from four-day-old seedlings grown under **C)** constant white light (24h) or **D)** diurnal cycling white light (6h white light / 18h dark) conditions harvested 1h after subjective dawn. Shown are mean expression values (+/- SE, n = 3 biological replicates). Statistical analyses in B, C, and D were performed using a one-way ANOVA followed by a Tukey HSD test. In B, letters indicate statistically significant differences between samples (*P* ≤ 0.05). In C and D, statistical significance in respect to *fc2* is indicated as follows: n.s. = p-value ≥ 0.05, * = p-value ≤ 0.05, ** = p-value ≤ 0.01, *** = p-value ≤ 0.001. Closed circles represent individual data points.

We then tested if *cry2* affects canonical ^1^O_2_ retrograde signaling to control nuclear gene expression. To this end, RNA was extracted from four-day-old seedlings gown in 24h constant or cycling light conditions one hour post dawn. We have previously demonstrated that this time point effectively captures the nuclear response to the chloroplast ^1^O_2_ signal [22, 25]. Using RT-qPCR, we monitored steady state transcript of six genes induced by ^1^O_2_ retrograde signals. *SIB1* and *HSP26.5* were identified as ^1^O_2_ maker genes in *fc2* [20]. *BAP1* and *ATPase* were identified as ^1^O_2_ maker genes in *flu* mutants [18]. *ZAT12* and *CYC8* are general stress marker genes also controlled by the ^1^O_2_ signal in *fc2* [44]. Under cycling light conditions, all six marker genes were induced (only *ATPase* was not statistically significant) in *fc2* compared to wt (**Fig. 3c**). Surprisingly, neither *cry1* nor *cry2* had a large impact of this induction. *HSP26.5* was slightly reduced in *fc2 cry1* and *fc2 cry2* compared to *fc2*, but was still 188-and 234-fold induced in *fc2 cry1* and *fc2 cry2*, respectively, compared to wt. Furthermore, *ATPase* and *CYC8* were induced even further in *fc2 cry2* compared to *fc2*. We also assessed transcript levels under permissive constant light conditions. Under these conditions, only *HSP26.5* was significantly induced in *fc2* compared to wt, albeit to a mild degree compared to its expression under cycling light conditions (3-fold vs. 362-fold) (**Fig. 3d**). Surprisingly, *cry2* led to a mild constitutive induction of *HSP26.5*, *ATPase*, *ZAT12*, and *CYC8* compared to *fc2*. These levels were 1-2 orders of magnitude less than in cycling light conditions, suggesting that the induction observed under cycling light conditions was dependent on bulk ^1^O_2_ accumulation. Together, these results suggest that *cry2* does not significantly affect canonical ^1^O_2_ retrograde signaling in *fc2*. Furthermore, it may lead to a mild constitutive stress response under permissive conditions.

### Global transcript analysis of *cry2* mutants

To follow-up on the above results, we performed an RNA-seq analyses on wt, *fc2*, *cry2*, and *fc2 cry2* seedlings to assess global patterns in steady-state transcripts. A Principal Component Analysis (PCA) analysis and distance matrix plot (**Figs. S6a and b**) were performed with the replicate samples (three each for a total of 24), with the largest principal component (PC1) corresponding to light conditions and PC2 corresponding to the presence or absence of *FC2*. Interestingly, global gene expression profiles in the wt and *cry2* backgrounds clustered together and were not clearly distinguishable. Similarly, *fc2* and *fc2 cry2* backgrounds clustered together and were not clearly distinguishable under cycling light conditions.

To identify genes controlled by ^1^O_2_ and/or CRY2, we performed four pairwise comparisons between wt and the mutants in permissive 24h light or ^1^O_2_-inducing cycling light conditions. We then applied cutoffs (Log_2_ Fold-Change (FC) ≥ |1.0|, *padj* ≤ 0.05) to identify differentially expressed genes (DEGs) between groups. Under constant light conditions, 541 differentially expressed genes (DEGs) were identified in *fc2* compared to wt, with 228 and 313 DEGs up-regulated or down-regulated in *fc2*, respectively (**Table S1, Figs. 4a and b**). A gene ontology (GO)-term enrichment analysis of the up-regulated genes shows that they are broadly involved in abiotic stress (enriched terms included: response to H_2_O_2_, heat, hypoxia, oxidative stress, and salt stress) (**Table S2**, **Fig. S7a**). The down-regulated genes were enriched for genes broadly involved in metabolism, including the terms photosynthesis light harvesting in photosystem II (PSII) and photosystem I (PSI), nicotinamide biosynthesis, and glucosinolate biosynthesis (**Table S2**, **Fig. S7b**). This pattern is consistent with *fc2* mutants accumulating less chlorophyll than wt and exhibiting reduced growth [20, 39].

**Figure 4.**
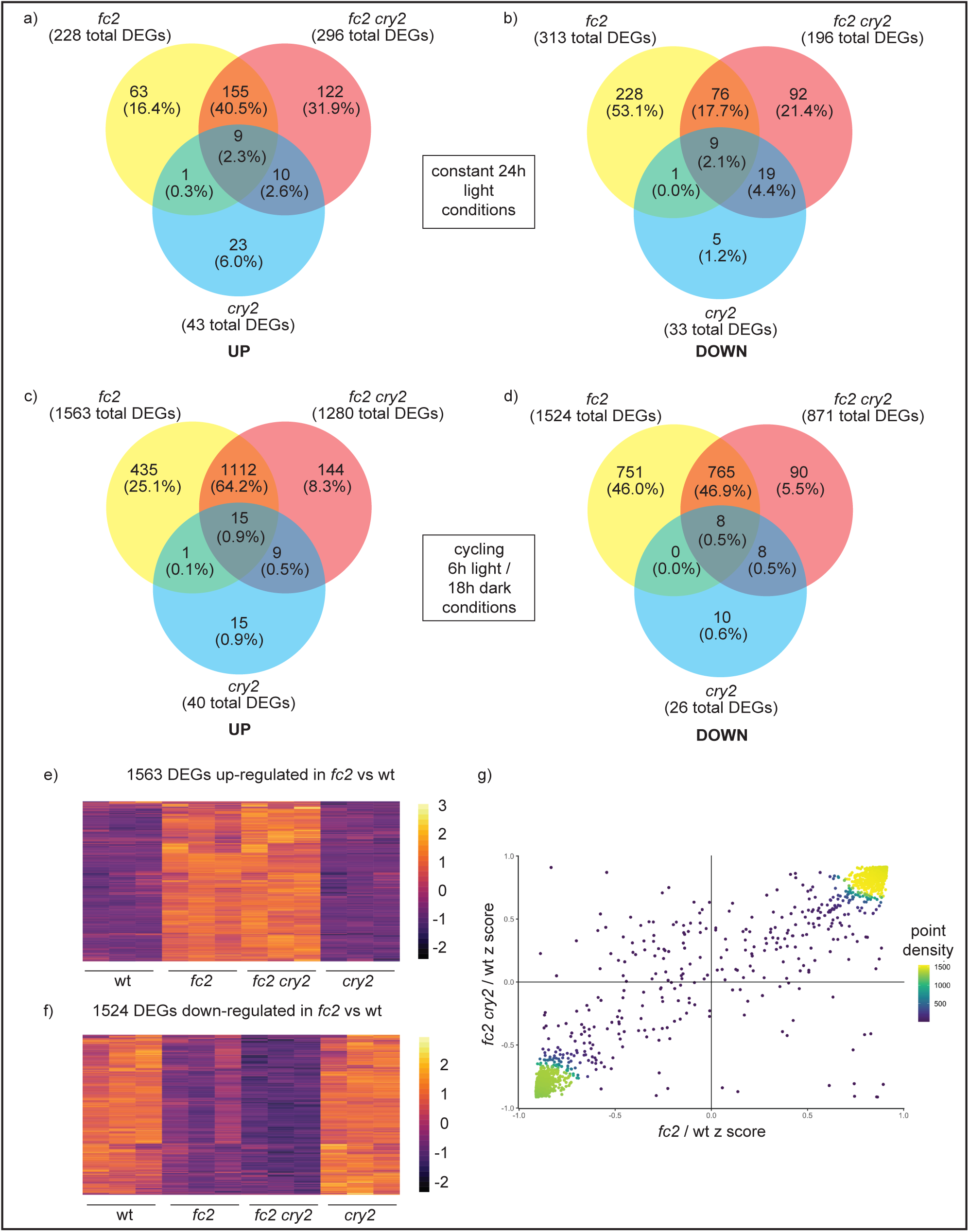
The global transcriptional response to singlet oxygen is not largely affected by the *cry2* mutation. An RNA-Seq analysis was performed to measure steady-state transcript levels in plants grown in permissive constant (24h) light conditions or singlet oxygen (^1^O_2_)-producing 6h light / 18h dark diurnal (cycling) light conditions for four days. Differentially expressed genes (DEGs) were identified by pairwise comparisons between genotypes in one condition (Log_2_FC ≥ +/- 1.0, P*adj* ≤ 0.05). **A-D)** Venn diagrams depicting unique and shared up-regulated and down-regulated DEGs in 24h light and cycling light conditions. **E** and **F**) Expression heatmap of the 1563 and 1524 DEGs up-regulated or down-regulated, respectively, in *fc2* vs. wt in cycling light conditions. The z-score color bar at right of each image shows the range of expression values, from decreased expression (purple) to increased expression (orange). Genes are ordered based on Euclidian distance clustering implemented as the default approach in the Pheatmap R package version 1.0.3 **G)** Scatter plot showing gene expression change relative to Col-0 in *fc2 cry2* double mutant (y-axis) or *fc2* alone (x-axis). Points are colored based on their neighboring density to allow for overlapping point estimation. The distance for two points to be considered neighbors was estimated using the ggpointdensity R package version 0.1.0 using default parameters.

Under cycling light conditions, the number of DEGs between *fc2* and wt increased: 1563 and 1524 genes were up- and down-regulated in *fc2*, respectively (**Figs. 4c and d**, **Table S3**). Among the up-regulated genes, there was a large overlap with those induced in constant light conditions (108 genes, 47%). As in constant light conditions, a GO-term enrichment also found terms related to abiotic stress (cellular response to hypoxia, abiotic stimulus), but also biotic stress (response to other organism, defense response) (**Fig. S8a**, **Table S4**). Among genes repressed in *fc2* cycling light conditions, there was less overlap with those found in constant light conditions (51 genes, 16%). Most of the enriched GO-terms were still related to photosynthesis light reactions, but also included the carbon reactions (**Fig. S8b**, **Table S4**.). This is consistent with earlier reports that chloroplast ^1^O_2_ down-regulates PhANGs [21, 45]. Our previous study [27] used a meta-analysis to identify a core set of 36 (25 up-regulated and 11 down-regulated) genes that respond to chloroplast ^1^O_2_ signaling. Among this core set, 23 of the 25 up-regulated genes and 9 of the 11 down-regulated genes were among the identified DEGs in *fc2* grown in cycling light. In constant light however, only 2 of the 25 up-regulated genes and 1 of the 11 down-regulated genes were in the *fc2* data set. Thus, the core set of genes are likely only activated under severe ^1^O_2_ stress and/or when cellular degradation is activated.

Consistent with our RT-qPCR analyses, the *cry2* mutation had only a modest impact on the transcriptional response in *fc2*, particularly for genes that are up-regulated. Compared to wt under constant light conditions, *fc2 cry2* mutants still up- and down-regulate 164 (72%) and 85 (27%) of the DEGs identified in *fc2*, respectively (**Figs. 4a and b, Table S5**). A GO-term enrichment of the shared up-regulated DEGs (i.e., those not reversed by the *cry2* mutation) was very similar to those identified in *fc2* vs wt and included terms generally associated with abiotic stress (**Fig. S9a, Table S6**). There were not enough up-regulated DEGs specific to *fc2* (i.e., not reversed by *cry2*) for a GO-term analysis.

Under cycling light conditions, a similar pattern was observed. A heatmap analysis of DEGs identified under cycling light conditions also shows that *fc2* and *fc2 cry2* are largely similar in their response (**Figs 4e and f**, **Table S7).** Compared to wt in these conditions, *fc2 cry2* mutants still share 1127 (72%) of up-regulated DEGs and 773 (51%) of down-regulated DEGs with *fc2* (**Figs. 4c and d**). A GO-term enrichment showed that up-regulated DEGs shared by *fc2* and *fc2 cry2* (i.e., not reversed by *cry2*) still included terms generally associated with abiotic and biotic stress (**Fig. S10a**, **Table S8**). However, up-regulated DEGs specific to *fc2* (i.e., reversed by *cry2*) did not include terms associated with abiotic stress, but those associated more broadly with metabolism (e.g., carboxylic acid, organic acid catabolic process) or JA responses, suggesting that *cry2* is not reversing abiotic stress responses in *fc2* in these conditions (**Fig. S10b, Table S9**). Down-regulated DEGs shared by *fc2* and *fc2 cry2* (i.e., not reversed by *cry2*) were similar to those in *fc2* and enriched for terms associated with photosynthesis (both light and carbon reactions) (**Fig. S10c, Table S8**). DEGs specific to *fc2* (i.e., reversed in *fc2 cry2*) were also enriched for photosynthesis terms (**Table S9**). Among the core ^1^O_2_-responsive genes [27], *cry2* had little effect on their expression (22 of 25 of the up-regulated genes and 8 of 11 of the down-regulated genes were still differentially expressed in *fc2 cry2* compared to wt).

The above analyses suggest that *cry2* does not block ^1^O_2_-induced cellular degradation by blocking the ^1^O_2_-induced retrograde signal for nuclear gene expression. Therefore, we investigated if the loss of CRY2 was causing unique changes to the transcriptome that were leading to reduced ^1^O_2_-induced PCD. To this end, we identified DEGs in *fc2 cry2* vs. wt that were not present in *fc2* vs. wt. In constant light conditions, 132 and 111 DEGs were identified that were induced or repressed, respectively, in *fc2 cry2* (but not *fc2*) compared to wt (**Figs. 4a and b**). The up-regulated genes were enriched for terms associated with abiotic stress (response to hypoxia, stress, and abiotic stimulus). Along with the previous RT-qPCR results, this indicated that *fc2 cry2* mutants may be experiencing constitutive stress even under permissive conditions (**Fig. S11a**, **Table S10**). In support of this, we identified 48 DEGs shared between *fc2 cry2* vs. wt in 24h conditions and *fc2* vs. wt *in* cycling light conditions, but not in *fc2* vs. wt in 24h light conditions (i.e., genes induced by ^1^O_2_, but constitutively expressed in *fc2 cry2*). These DEGs were enriched for terms related to hypoxia stress (**Fig. S11b**, **Table S11b**), suggesting that *fc2 cry2* may have a higher basal expression of ^1^O_2_-controlled stress genes, which may prime *fc2 cry2* mutants for photo-oxidative stress. These differences were much larger than with the *cry2* mutation alone. In constant light conditions, we only identified 43 and 33 up-regulated and down-regulated DEGs (**Table S12**), respectively, in *cry2* vs. wt, which were not enriched for genes involved in stress responses (**Fig. S11c, Table S13**).

To further test if *cry2* was influencing gene expression in the *fc2* background, we next directly compared transcript levels between *fc2* with *fc2 cry2* in cycling light conditions. Under cycling light conditions, only 65 and 27 up-regulated and down-regulated DEGs were identified, respectively (**Table S14**), which is only 3% of the response observed in *fc2* vs wt. This included only 2 of the 36 ^1^O_2_ core response genes. A GO-term enrichment analysis did not reveal any significantly enriched terms for the up-regulated genes, but genes broadly related to defense and biotic stress were enriched in the down-regulated DEGs (**Table S15**, **Fig. S11d**), indicating that some of the stress response has been dampened by *cry2*, possibly indirectly due to an absence of cellular degradation. We next performed an opposing expression analysis to determine which genes were both induced in *fc2 cry2* and repressed in *fc2* (or vice versa) (**Fig. 4g**). As predicted by the analysis of shared DEGs, relatively few genes were found to be opposingly expressed, with 54 and 22 genes up-regulated and down-regulated in *fc2 cry2*, respectively (**Table S16**). No GO-terms were significantly enriched in the down-regulated genes, but the up-regulated genes were enriched with terms related to photosynthesis and chloroplast function (**Table S17**). 30 of the genes (56%) were plastid-encoded genes, 8 of which encode components of the NADH:ubiquinone oxidoreductase involved in cyclic electron flow. Such an enrichment of plastid transcripts in *fc2 cry2* may be due to the decreased chloroplast degradation rates in this mutant.

Together, the global transcript profiles suggest that the *cry2* mutation does not block PCD and chloroplast degradation by reducing ^1^O_2_ or canonical retrograde signaling to the nucleus. Instead, it may lead to the constitutive expression of stress-related genes under mild photo-oxidative stress, thereby acclimating plants to high levels of ^1^O_2_.

### CRY2 controls singlet oxygen-induced cellular degradation independent of blue light signaling

The finding that CRY2 is involved in propagating the ^1^O_2_ signal to control cellular degradation, suggests that blue light signaling by CRY2 may be part of the signaling mechanism. To test this possibility, we first determined if blue light is required for the *fc2* PCD phenotype. To this end, we grew seedlings under light cycling conditions for six days, but used monochromatic blue (bc) or red light (rc) instead of white light (wc). Using a titration of light intensities, we determined that 75 µmol photons m^-2^ sec^-1^ bc was sufficient to induce PCD in *fc2* (**Fig. S12a**). Next, we performed a similar titration of rc and found that 40 µmol photons m^-2^ sec^-1^ rc was sufficient to induce PCD in *fc2* (**Fig. S12b**), suggesting that bc is not strictly required for the *fc2* PCD phenotype.

Next, we hypothesized that if CRY2 requires bc to activate the *fc2* PCD phenotype, *fc2 cry2* mutants should resemble *fc2* mutants under cycling rc conditions. To test this, we grew seedlings under cycling rc conditions and assessed PCD after six days. Both *fc2 cry2* and *fc2 pub4* retained their ability to survive under these conditions and exhibited significantly less PCD that *fc2* (**Fig. 5a**). A trypan blue stain confirmed these results (**Figs. 5b and c**)). This suggests that, even in the absence of bc, CRY2 is necessary for ^1^O_2_-induced PCD in *fc2*. We also tested phenotypes under cycling bc conditions and observed the same pattern of PCD among the genotypes (**Figs. 5d-f**). Furthermore, *fc2 cry1* still experienced PCD, further confirming that CRY1 and cryptochrome-dependent bc signaling is not required for ^1^O_2_-induced PCD in *fc2* [27].

**Figure 5.**
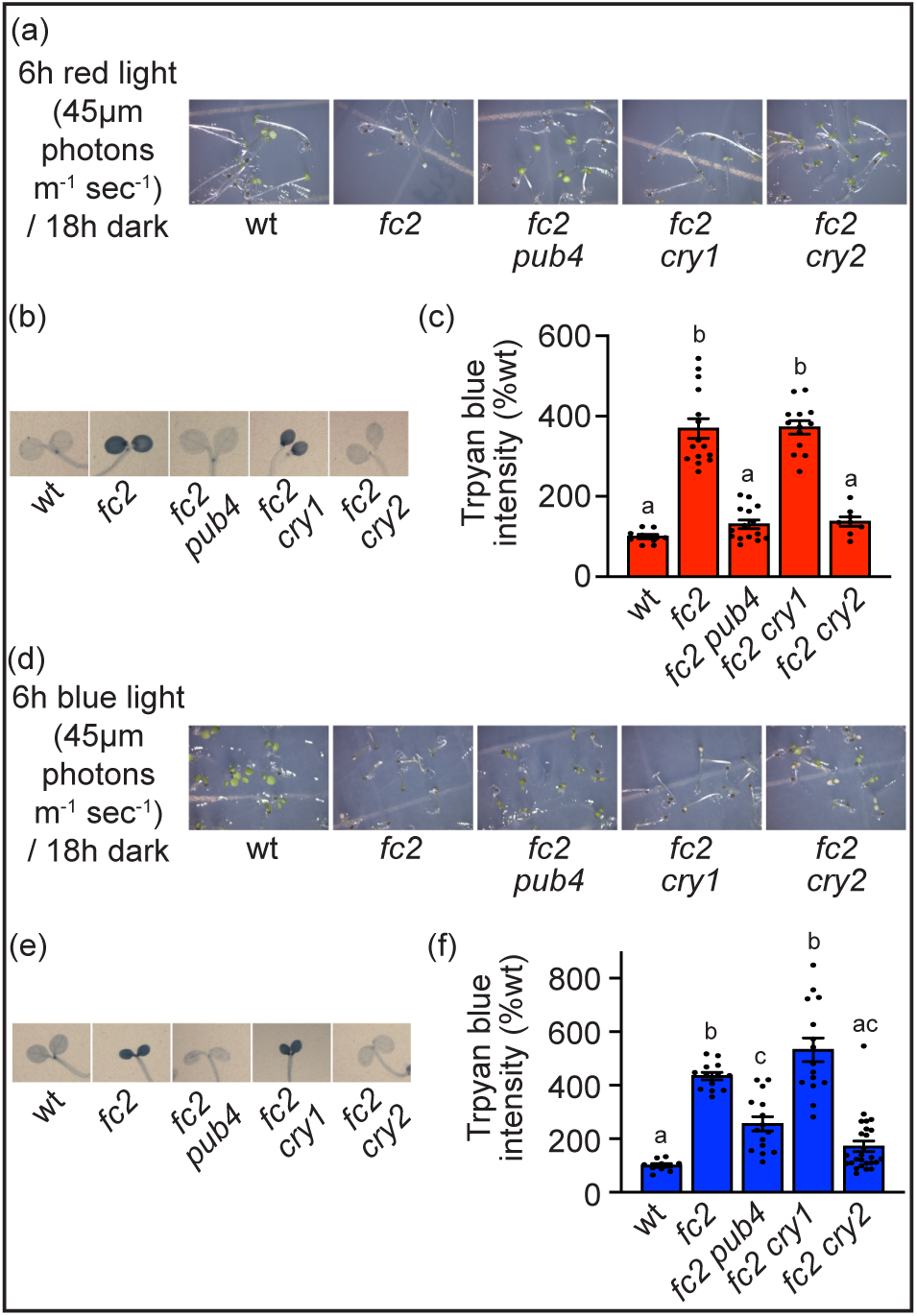
The *fc2* cell death phenotype can be induced independent of blue light signaling. The *fc2* cell death phenotype was tested under monochromatic light conditions. **A)** Shown are six-day-old seedlings grown under diurnal cycling red light (6h red light [45 µmol photons m^-2^ sec^-1^] / 18h dark) conditions. **B)** Representative seedlings from panel A stained with trypan blue. The dark blue color is indicative of cell death. **C)** Mean intensities of the trypan blue signal in B (+/- SE, n ≥ 4 seedlings). **D)** Shown are six-day-old seedlings grown under diurnal cycling blue light (6h red light [65 µmol photons m^-2^ sec^-1^] / 18h dark) conditions. **E)** Representative seedlings from panel D stained with trypan blue. The dark blue color is indicative of cell death. **F)** Mean intensities of the trypan blue signal in E (+/- SE, n ≥ 4 seedlings). Statistical analyses in **C** and **F** were performed using a one-way ANOVA followed by a Tukey HSD test. Letters indicate statistically significant differences between samples (*P* ≤ 0.05). Closed circles represent individual data points.

The above results suggest that CRY2-mediated blue light signaling is not required for PCD in the *fc2* mutant and that CRY2 may be acting through a different mechanism. To further test this hypothesis, we complemented *fc2 cry2* with three versions of *GFP-CRY2* driven by the constitutive *ACT2* promoter: wt *CRY2* (*ACT2*::*GFP-CRY2*), a mutant CRY2 unable to bind FAD and be photo-activated by bc (*ACT2*::*GFP-CRY2*-D387A) [46], and a mutant CRY2 unable to interact with COP1 [47] and SPA1 [48] and promote canonical bc-dependent signaling [32] (*ACT2*::*GFP-CRY2*-P532L). We also transformed plants with *ACT2*::*GFP* as a negative control. Next, we tested the ability of these constructs to complement the *cry2* blue light signaling defect. Seedlings were grown for five days under 1 µm photons bc m^-2^ sec^-1^ and hypocotyl elongation was scored. *fc2 cry2* seedlings (and those transformed with *ACT2*::*GFP*) had significantly elongated hypocotyls, which was fully complemented by wt *GFP*-*CRY2* (**Fig. 6a**). As expected, neither mutant form of *CRY2* was able to complement the elongated hypocotyl phenotype due to these proteins’ bc signaling functions being compromised [32]. Next, we tested the phenotypes of these lines under light cycling conditions to induce ^1^O_2_ production. As expected, wt *GFP-CRY2* complemented the *fc2 cry2* phenotype and restored PCD (**Fig. 6b**). Unexpectantly, both mutant forms of GFP-CRY2 were also able to restore PCD to the *fc2 cry2* mutant. A trypan blue stain confirmed these phenotypes (**Figs. 6c and d**). Together, these results further support the conclusion that the bc signaling activity of CRY2 is not necessary for its ^1^O_2_ signaling function.

**Figure 6.**
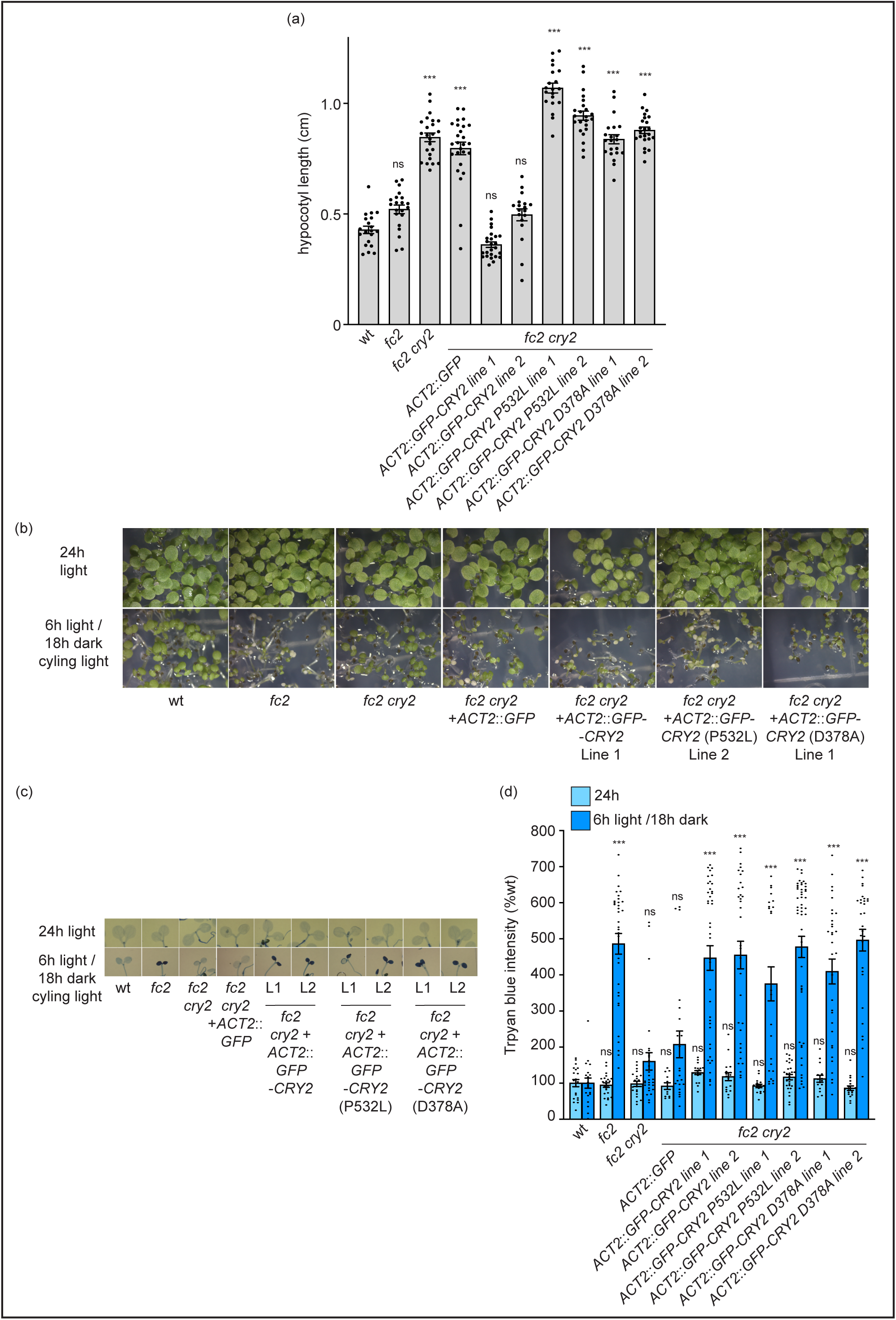
CRY2 does not propagate singlet oxygen signaling via canonical blue light signaling mechanisms. *cry2* mutants lacking blue light signaling activity were tested for their ability to complement *fc2 cry2* phenotypes. **A)** Shown are the mean hypocotyl lengths of five-day-old seedlings grown under constant blue light (1 µmol photons m^-2^ sec^-1^) conditions (+/- SE, n ≥ 18 seedlings). **B)** Shown are six-day-old seedlings grown under 24h constant light or diurnal light cycling (6h light / 18h dark) conditions. **C)** Representative images of seedlings from panel A stained with trypan blue. The dark blue color is indicative of cell death. **D)** Shown is the quantification of trypan blue stain of the trypan blue signal in C (+/- SE, n ≥ 12 seedlings). Statistical analyses in **A** and **D** were performed using a one-way ANOVA followed by a Tukey HSD test. Statistical significance in respect to *fc2* is indicated as follows: n.s. = p-value ≥ 0.05, * = p-value ≤ 0.05, ** = p-value ≤ 0.01, *** = p-value ≤ 0.001. Closed circles represent individual data points.

Another study recently demonstrated that CRY2 has important signaling functions independent of bc, and 834 genes were shown to be induced in etiolated *cry2* seedlings grown in the dark [49]. To test if any of these CRY2-dependent/bc-independent (CD/BI) genes are being regulated by CRY2 in our experiments, we compared them to CRY2-related DEGs identified in our study. If we utilize the cutoffs used in the previous study (fold change ≥ |1.5|, *padj* ≤ 0.05), 82 genes are up-regulated in *cry2* vs wt in 24h light (**Tables S18**). 13 significantly overlap with CD/BI DEGs (hypergeometric analysis = 9.99E-08) and are enriched for photosynthesis (six genes encode for Lhcb/Cab proteins involved in light harvesting in the chloroplast) (**Fig. S13a**). This suggests that at least some CD/BI DEGs can be regulated in the light to affect ^1^O_2_ signaling in our conditions.

The 834 CD/BI DEGs previously identified in etiolated seedlings were enriched for GO terms related to “photosynthesis”, “high light” “decreased oxygen” and “response to stress” (**Fig. S13b**) and are similar to those found to response to ^1^O_2_ (**Fig. S8a**). To determine of any of the CD/BI DEGs are involved in ^1^O_2_ stress, we compared genes induced in *fc2* compared to wt in cycling light conditions. 76 significantly overlap (hypergeometric test = 1.15E-08) (**Table S18**) and were enriched for terms related to “hypoxia”, “abiotic” and “biotic stress,” further suggesting that *cry2* leads to a transcriptional priming to ^1^O_2_ stress (**Fig. S13c**). Together, this further points to a blue light-independent role for CRY2 in promoting a ^1^O_2_ signal to induce cellular degradation, possibly by repressing the expression of genes involved in stress and photosynthesis.

## Discussion

To further understand the molecular mechanisms behind chloroplast ^1^O_2_ signaling, we systematically tested if photoreceptors play a role in promoting these signals in *fc2* mutants. Of the mutants tested, *cry2* robustly blocked ^1^O_2_-induced cellular degradation, including PCD and chloroplast degradation. ^1^O_2_ still accumulates in *fc2 cry2* mutants, suggesting that a signal, rather than photo-oxidative stress, is being blocked in this mutant. Importantly, the impact of ^1^O_2_ on retrograde signaling and nuclear gene expression was largely unaffected and transcript profiles of *fc2* and *fc2 cry2* are remarkably similar when grown under cycling light conditions (**Figs. 4e and f**), demonstrating that the two main cellular responses to ^1^O_2_ (retrograde signaling and cellular degradation) can be uncoupled. This makes *cry2* unique as other genetic suppressors of ^1^O_2_-induced cellular degradation in *fc2* have been shown to strongly mitigate retrograde signaling. An analysis of ^1^O_2_ marker genes in *fts* mutants affecting plastid gene expression (*ppr30*, *mterf9*, *ctps2*) and an RNA-seq analysis of an *fts* mutant that affects chloroplast ubiquitination (*pub4*) demonstrated that these mutations mostly return nuclear gene expression (both induced and repressed) to wt levels [21, 22, 25]. This suggests that CRY2 likely functions further downstream of chloroplast stress, only affecting part of the cellular response. This is in line with our observations that the *cry2* mutation does not strongly affect chloroplast development or plastid gene expression, and that it must be blocking ^1^O_2_ signaling through a mechanism independent of chloroplast function. The nuclear localization of CRY2 further implicates it as acting downstream of other known ^1^O_2_ signaling factors, possibly at a point where signals for retrograde signaling and cellular degradation diverge.

This uncoupling of phenotypes also suggests that the bulk of ^1^O_2_-controlled gene expression is not a response to cellular degradation, but an acclimation response to photo-oxidative stress. A similar conclusion was drawn from the ability of *cry1* to block ^1^O_2_-induced PCD in the *flu* mutant without affecting ^1^O_2_ levels [35]. In that study, a microarray analysis showed that *cry1* had a limited impact on nuclear gene expression. The induction of only 34 (6%) ^1^O_2_-responsive genes was blocked by *cry1,* and these were hypothesized to be necessary for initiating PCD. This further illustrates that the impact of ^1^O_2_ on retrograde signaling and cellular degradation can be separated, but the roles of CRY1 and CRY2 in signaling appear to be different. First, CRY1 was previously shown to be dispensable for ^1^O_2_-induced PCD and retrograde signaling in *fc2* [27], while the impact of *cry2* on ^1^O_2_-induced PCD in *flu* was negligible [35]. Second, the role of CRY1 in ^1^O_2_ signaling is blue light-dependent [35], whereas the role of CRY2 in ^1^O_2_ signaling is independent of blue light activation (**Figs. 5a-c** and **6b-d**). Third, CRY1 was shown to regulate the expression of 34 genes that were only induced six hours after ^1^O_2_ accumulation [35]. In *fc2*, 15 of these genes are already induced within one hour of ^1^O_2_ stress, and only two of these were reversed by *cry2* (**Table S19**). Together, these results reinforce the hypothesis that multiple ^1^O_2_ signals likely exist in *fc2* and *flu* [27], and the cryptochromes may be playing different signaling roles in each.

Our results support the conclusion that the role of CRY2 in promoting the ^1^O_2_ signal is not dependent on its blue light activation and signaling. First, cycling monochromatic red light was sufficient to induce PCD in *fc2*, in a CRY2-dependent manner (**Figs. 5a-c**). Second, PCD in *fc2 cry2* could be restored by expressing mutant forms of CRY2 that lacked FAD chromophore binding (D378A) or the ability to interact with COP1 and activate canonical blue-light signaling capabilities (P532L) (**Figs. 6b-d**). As expected due to impaired blue light signaling, these constructs failed to complement blue-light inhibition of hypocotyl growth (**Figs. 6a**) [32], demonstrating that the ability to propagate the ^1^O_2_ signal is independent from these canonical functions.

These results were surprising as most functions of CRY2 have been attributed to its blue light activation, and subsequent oligomerization and interactions with COP1 and other signaling proteins [32]. However, it has recently been reported that CRY2 plays an important, and long overlooked, role in dark-grown etiolated seedlings [49], where 1,534 genes are misregulated in *cry2* mutants compared to wt. This reprogramming affected root cell division, suggesting that CRY2 at least impacts root development independent of light. Interestingly, many of the genes mis-regulated in etiolated *cry2* seedlings are associated with chloroplast function and photosynthesis (**Fig. S13b**) and ^1^O_2_ stress signaling (**Fig. S13c**), suggesting that CRY2 also plays an important blue light-independent role in shoots, possibly involving chloroplast function or signaling. For a subset of these genes, this regulation by CRY2 is dark-dependent and reversed after 30 minutes of blue light irradiance [49]. Overall, this could allow for the coordination of chloroplast development with light to minimize photo-oxidative damage, which can lead to premature seedling death before photomorphogenesis is completed [50]. In *fc2*, however, the role of CRY2 does not appear to be strictly dark-dependent. In *fc2 cry2* mutants, PCD is blocked after light exposure (**Figs. 1a, c, and d**) and chloroplast degradation is blocked under constant light conditions (**Figs. 2a and b**). Together, our results expand on the blue light-independent role of CRY2 and show that it is also important in photosynthetic tissue, possibly to regulate the response to photo-oxidative stress from developing chloroplasts.

How CRY2 may be acting in a blue light-independent manner to control ^1^O_2_ signaling is not clear. In roots, non-photoactivated CRY2 physically interacts with FORKED-LIKE 1 (FL1) and FL3 to control root division genes via chromatin remodeling [49]. Whether these interactions occur in the shoot, or if non-photoactivated CRY2 interacts with different proteins to regulate gene expression is not known. A different mechanism may be likely, as the role of CRY2 in ^1^O_2_ signaling is not dark-dependent and can occur in the light. In the shoot, CRY2 may signal through a different set of interacting proteins, or possibly involve interactions with other photoreceptors. For instance, CRY2 can inhibit hypocotyl elongation in red light [51] and CRY1 and CRY2 have been shown to interact with phyA and phyB, respectively [51, 52]. This may allow cryptochromes to modulate responses to red and/or far-red by affecting these photoreceptors and downstream PIF function [31]. Indeed, our results show that phyA does play a stage-specific role in modulating PCD in response to ^1^O_2_ accumulation (**Figs. 1e-g**). Whether CRY2 and phyA function together to regulate ^1^O_2_ signaling will need to be further investigated.

How CRY2 regulates PCD in response to chloroplast ^1^O_2_ is also not entirely clear. Our RT-qPCR analysis of chloroplast ^1^O_2_ stress marker genes showed that they are constitutively expressed under permissive constant light conditions in *fc2 cry2* mutants (**Fig. 3d**). This is different from other *fts* mutants, where these genes are not affected (*pub4*) [20] or mildly repressed (*ppr30*, *mterf9*, *ctps2*) [22, 25]. The RNA-seq analysis also demonstrated that *fc2 cry2* constitutively up-regulates DEGS enriched for stress terms even under permissive constant light conditions; 48 DEGs that are induced by ^1^O_2_ in *fc2* (**Fig. S11b**) and 132 unique DEGs not induced *fc2* (**Fig. S11a**). As such, *cry2* may transcriptionally prime cells for acclimation to ^1^O_2_ stress, which dampens the activation of PCD and chloroplast degradation without repressing retrograde signaling. The opposing expression analysis of *fc2* vs. *fc2 cry2* under cycling light conditions (**Fig. 4g**), showed an enrichment of plastid-encoded transcripts in *fc2 cry2*, which may indicate that levels of chloroplast degradation is reduced in *fc2 cry2*. However, as the RNA-seq libraries were enriched for polyadenylated transcripts, it may be that plastid transcripts in *fc2 cry2* are polyadenylated and targeted for turnover [53, 54], indicating that *fc2 cry2* chloroplasts may be undergoing a major programmatic shift to tolerate photo-oxidative damage. In any case, this heightened state of stress response in the *fc2 cry2* mutant at least partly depends on the *fc2* background, possibly due to the low levels of chloroplast ^1^O_2_ responsible for selective chloroplast degradation [24]. Interestingly, plant cryptochromes have been shown to generate O_2_^-^ (and subsequently H_2_O_2_) via reoxidation of FAD by molecular oxygen, which can alter gene expression in Arabidopsis cells [55, 56]. Such a function may allow plants cells to acclimate to photo-oxidative stress, although this ability of CRYs appears to be dependent on chromophore binding and is likely blue-light dependent [56]. Nonetheless, the possibility that CRYs can generate ROS, either directly or indirectly, in a blue light-independent manner has not been thoroughly tested.

It is also possible that CRY2 may control a small subset of ^1^O_2_-induced genes that promote or inhibit cellular degradation. As discussed above, a similar conclusion was drawn for the role of CRY1 in regulating ^1^O_2_-induced PCD in the *flu* mutant [35]. Here, *cry2* had a relatively minor impact on transcript abundance in the *fc2* background and the number of DEGs identified between *fc2* and *fc2 cry2* was 3% of those identified between *fc2* and wt, but was enriched for terms related to defense and biotic stress (**Fig. S11d**). As in *flu*, it is also possible that one hour is not sufficient to induce such cellular degradation-associated genes. PCD in *fc2* is not observed until at least three hours after light exposure [20] and such a transcriptional response may have been missed by our analysis. Alternatively, CRY2 may regulate gene expression in response to ^1^O_2_ via a post-transcriptional mechanism. This is an intriguing possibility as CRY2 has recently been shown to regulate the alternative splicing of at least some transcripts (e.g., *FLOWERING LOCUS T* [*FT*]) [57]. However, at least in that reported case, regulation requires the blue-light activation of CRY2, and it is unknown if CRY2 can regulate post-transcriptional regulation through blue light-independent mechanisms.

Together, our results demonstrate that CRY2 plays a previously unknown blue light-independent role in photosynthetic tissue to promote the induction of PCD and chloroplast degradation in response to chloroplast ^1^O_2_ and photo-oxidative stress. CRY2 is unique in that it is the only ^1^O_2_ signaling factor in *fc2* shown to not strongly impact retrograde signaling, suggesting that the ^1^O_2_ signal diverges after leaving the chloroplast and controls cellular degradation and nuclear gene expression separately. How CRY2 impacts cellular degradation from its nuclear localization is unknown, but we hypothesize that it may involve the transcriptional priming of cells for ^1^O_2_ stress to help plants coordinate cellular degradation in response to severe stress. The further investigation of how CRY2 regulates gene expression independent of blue light, should help to reveal the mechanisms by which this is achieved and how CRYs control plant development under dynamic environments.

## Materials and Methods

### Biological material, growth conditions, and treatments

The *Arabidopsis thaliana* ecotype Columbia (Col-0) was used as wt and the genetic background except where the ecotype Landsberg erecta (Ler) is indicated. The *fc2-1* T-DNA insertion line (GABI_766H08) is from the GABI collections [58] and was previously described [59]. The *cry1-304* [60] *cry2-1* [37], *phyA-211* [61], *phyB-9* [62], and *pub4-6* [20] mutants were previously described. The *cry2* alleles *fha-1*, *fha-2*, and *fha-3* in the Ler background were previously described [63]. Mutant lines used in this study are listed in **Table S20**. Single mutants were crossed to produce double and triple mutant combinations, and all mutants were confirmed by PCR analyses (primer sequences listed in **Table S21**). The *fc2-1* mutation was introgressed into the Ler background by outcrossing nine times, confirming the presence of the *fc2-1* T-DNA by PCR in each generation.

Seeds were surface sterilized and plated in two ways as previously described [25]. 1) Seeds were washed in 30% bleach solution (v:v) with 0.04% Triton X-100 (v:v) in 10 minutes and then rinsed with sterile water three times by pelleting seeds at 2,000 x g for 30 seconds. 2) 25-100 µl of seed was placed in 2 mL microcentrifuge tubes within an airtight chamber with their lids open. A beaker of 150 mL of bleach was mixed with 5 mL of concentrated HCl and added to the chamber before sealing. Seeds were removed 16-24h later and plated. Sterilization Method 1 was used for plants monitored or assayed in the seedling stage. Sterilization Method 2 was used for growing plants for bulking seed, genotyping, and adult-stage experiments. Seeds were plated on Linsmaier and Skoog medium pH 5.7 (Caisson Laboratories North Logan, UT) with 0.6% micropropagation type-1 agar powder. Seeds were stratified for 3 to 5 days in the dark at 4°C prior to germination. Unless otherwise specified, standard conditions used to grow seedlings were ∼75 μmol photons m^-2^ sec^-1^ at 21°C in fluorescent light chambers (Percival model CU-36L5), set to control conditions (constant [24h] light) or stress conditions (cycling light [6h light/18h dark]). For adult stage experiments, 7-day-old seedlings were transferred to soil and grown in fluorescent reach-in growth chambers (Conviron), set to constant light or cycling light (16h light/8h dark) conditions. All above experiments were performed in chambers with cool white fluorescent bulbs. Light intensities were measured with a LI-250A light meter with a LI-190R-BNC-2 Quantum Sensor (LiCOR). Monochromatic light experiments were conducted in a Percival LED-30L1 growth chamber. Light spectra were measured using a PS-100 Spectroradiometer (Apogee Instruments).

All bacteria (*Escherichia coli* and *Agrobacterium tumefaciens* strains) were grown in liquid Miller nutrient broth or solid medium containing 1.5% agar (w/v). Cells were grown at 37°C (*E. coli*) or 28°C (*A. tumefaciens*) with the appropriate antibiotics and liquid medium was shaken at 225 r.p.m.

### *CRY2* complementation analyses

A wt copy of the *CRY2* cDNA was obtained by PCR amplifying from wt Arabidopsis cDNA DNA template using the Q5 enzyme (New England Biolabs). Primers WLO1904 and WLO1906 were used to amplify a 1,900 bp fragment containing the entire 1,839 bp coding region (start to sop codon) (Primers listed in **Table S21**). The DNA fragment was gel-purified using the Zymoclean Gel DNA Recovery Kits (Zymo) and cloned into the Gateway-compatible vector pDONR221 (Invitrogen) using the BP clonase kit (Invitrogen) according to the manufacturer’s instructions. The DNA fragment was then transferred to pEARLEYGATE101 (*35S* overexpression promoter [64]) to create pWLP396 (*35S*::*CRY2*) using the LR clonase kit (Invitrogen) according to the manufacturer’s instructions. *ACT2*::*GFP-CRY2* vectors are previously described [32]. All primers and vectors are listed in **Tables S21** and **S22**, respectively. Binary vectors were then transformed into *A. tumefaciens* strain GV301, which was subsequently used to transform Arabidopsis via the floral dip method. T1 plants were selected for their ability to grow on Basta-soaked soil and propagated to the next generation. T2 lines were monitored for single insertions (segregating 3:1 for Basta resistance:sensitivity) and propagated to the next generation. Finally, homozygous lines were selected in the T3 generation based on 100% Basta resistance.

### RNA extraction and real-time quantitative PCR

Total RNA extraction, cDNA synthesis, and RT-qPCR was performed following previously described protocols [25]. The RNeasy Plant Mini Kit (Qiagen) was used to extract total RNA from whole seedlings. Next, cDNA was synthesized using the Maxima first strand cDNA synthesis kit for RT-qPCR with DNase (Thermo Scientific) following the manufacturer’s instructions. Real-time PCR was performed using the SYBR Green Master Mix (BioRad) with the SYBR Green fluorophore and a CFX Connect Real Time PCR Detection System (BioRad). The following 2-step thermal profile was used in all RT-qPCR: 95 °C for 3 min, 40 cycles of 95 °C for 10s and 60 °C for 30s. *ACTIN2* expression was used as a standard to normalize all gene expression data. **Table S21** lists the primers used.

### Measuring bulk singlet oxygen levels

Bulk accumulation of ^1^O_2_ in seedling cotyledons was conducted as previously described [25] using Singlet Oxygen Sensor Green (SOSG, Molecular Probes) and imaged with a Zeiss Axiozoom 16 fluorescent stereo microscope equipped with a Hamamatsu Flash 4.0 camera. Average SOSG fluorescence per pix^2^ was quantified using ImageJ, choosing the brightest of the two cotyledons per seedling.

### Assessment of cell death

Cell death in seedling cotyledons and adult leaf tissue was assessed by staining tissue with trypan blue as previously described [25]. Average trypan blue intensity per pix^2^ was quantified using ImageJ, choosing the darkest blue of the two cotyledons per seedling.

### Light response assays

Ligh responses in seedlings were assessed by measuring hypocotyl length under monochromatic light conditions. Seed germination was first initiated under fluorescent white light lamps (120 µmol photons m^-2^ s^-1^) for two hours at 21°C. Seedlings were then grown for five days in 24h constant blue light (451nm), red light (650-670nm), or far-red light (720-740nm) conditions in a LED-30L1 growth chamber (Percival). Fluence rates of 1 µmol (blue), 12 µmol (red), and 8 µmol (far-red) photons m^-2^ s^-1^ were used to stimulate function of CRY2 [37], phyB [62], or phyA [61]. Seedlings were transferred to transparency sheets and scanned using an EPSON VE70 Photo scanner and measured using ImageJ.

### Measuring total chlorophyll content

Total chlorophyll content was measured in seven-day-old seedlings as previously described [22]. Chlorophyll was measured spectrophotometrically, and path corrections were calculated as previously described [65].

### Protochlorophyllide measurements

Protochlorophyllide was extracted from whole seedlings with 80% acetone and measured by fluorescence in a Varioskan LUX Microplate Reader using an adapted protocol [66] as described previously [20].

### RNA-Seq analyses

For whole transcriptome RNA-Seq analyses, seedlings were grown under 24h constant light or diurnal light cycling (6h light/18h dark) for four days and whole seedlings were collected one hour after subjective dawn. Samples were collected in biological triplicates, each replicate containing a pool of seedlings grown on a separate plate. Total RNA was extracted using the RNeasy Plant Mini Kit (Qiagen) and residual DNA was removed with the RNase-Free DNase Kit (Qiagen). A Qubit 2.0 Fluorometer was used to calculate RNA concentration and 4 mg of total RNA was used to prepare libraries with the TruSeq Stranded mRNA Library Prep Kit (Illumina, San Diego, US). Single-end sequencing (50bp) was performed on an Illumina HiSeq 2500.

### Identification of differentially expressed genes and gene expression analysis

Single-end 50bp reads were processed using the RMTA pipeline version 2.6.3 [67] using the HiSat2 v2.1.0 [68] and Stringtie v1.3.4 [69] mapping and assembly options. For expression quantification, reads were quantified against the *Arabidopsis thaliana* Araport11 transcriptome [70] using Salmon version 1.10.1 [71] against a decoy aware index, with the following arguments set: --gcBias, --posBias, --seqBias. Salmon quantification files were then loaded into R using the tximport version 1.28 package [72]. Differential expression was assessed using the DESeq2 package version 1.48.2 [73]. Genes with a Log2 fold-change > |1.0| and a Padj < 0.05 were considered differentially expressed genes (DEGs).

Gene Ontology (GO) enrichment analyses were performed using ShinyGO 0.80 (http://bioinformatics.sdstate.edu/go/). The significantly enriched GO terms were determined using default settings and a p-value threshold of ≤ 0.01.

### Alignment of *cry2* mutant sequences

Sequences of *cry2*/*fha* mutant candidates in the Ler *fc2* background were confirmed by amplifying the *CRY2* gene with Primers (WLO2046 and WLO2047, **Table S21**) and sequencing via the Sanger method (Eton Bioscience). The sequences were aligned to the *A. thaliana* TAIR10 reference genome using Geneious with default settings for nucleotide alignments.

### Transmission electron microscopy

For transmission electron microscopy (TEM), four-day-old seedlings grown in 24h constant light conditions were fixed, imaged, and analyzed as previously described [25]. Average chloroplast area was analyzed using ImageJ software. Only intact cells were used for these calculations. Measurements were collected from at least two cells from three different seedlings in each genotype.

## Declarations

### Data statement

Raw RNA-seq data and associated metadata files have been deposited at NCBI Gene Expression Omnibus and will be made publicly available upon acceptance of this manuscript. All other data is included in the manuscript or Supplementary Material. Further inquiries can be directed to the corresponding author.

### Authors’ contributions

DWT and JDW planned and designed the research. DWT created higher-order mutants and transgenic lines, performed genotyping, phenotyping, cloning, biochemical assays, RNA extractions, and cell death measurements. DTW and RAE performed RT-qPCR analyses. KEF performed all TEM analyses. KP and ADLN performed RNA-seq analyses. JDW conceived the original scope of the project and managed the project. DWT, RAE, KP, KEF, ADLN, and JDW contributed to data analysis and interpretation, and reviewed the manuscript. DWT and JDW wrote the manuscript.

## Acknowledgments

The authors thank Marta A. Kozlowksa (University of Arizona) for technical assistance with RNA-seq analyses, Dr. Matthew D. Lemke (University of Arizona) for technical assistance with the SOSG assays, and Dr. Chentao Lin (U. California, Los Angeles) for generously providing *FGFP-CRY2* complementation vectors.

## Supporting information

### Supplemental Figures

Figure S1. phyA and phyB are not required for singlet oxygen induced cell death in *fc2* seedlings.

Figure S2. Late flowering and suppression of cell death phenotypes are linked in *fc2 cry2*.

Figure S3. *cry2* mutant alleles suppress cell death in the Landsberg erecta *fc2* mutant.

Figure S4. Complementation of the *cry2* mutation.

Figure S5. Testing the impact of *cry2* on chloroplast development.

Figure S6. Visualization of variance among RNA-seq replicates used in the study.

Figure S7. Gene ontology analyses of differentially expressed genes in *fc2* vs. wt under constant light conditions.

Figure S8. Gene ontology analyses of differentially expressed genes in *fc2* vs. wt under diurnal cycling light conditions.

Figure S9. Gene ontology analyses of differentially expressed genes shared between *fc2* vs. wt and *fc2 cry2* vs. wt under constant light conditions.

Figure S10. Gene ontology analyses of differentially expressed genes identified in *fc2* vs. wt under cycling light conditions.

Figure S11. Gene ontology analyses of differentially expressed genes identified in *cry2* and *fc2 cry2* mutants.

Figure S12. Testing level of red and blue light necessary to induce cell death in *fc2* mutant seedlings.

Figure S13. Gene ontology analyses of blue light-independent differentially expressed genes identified in *cry2* and *fc2* mutants.

### Supplemental Tables

Table S1. All differentially expressed genes identified between *fc2* and wt under constant (24h) light conditions

Table S2. Gene Ontology term enrichment of DEGs identified between *fc2* and wt in constant light conditions

Table S3. All differentially expressed genes identified between *fc2* and wt under cycling (6h light / 18h dark) light conditions

Table S4. Gene Ontology term enrichment of DEGs identified between *fc2* and wt in cycling (6h light / 18h dark) light conditions

Table S5. All differentially expressed genes identified between *fc2 cry2* and wt under constant (24h) light conditions

Table S6. Gene Ontology term enrichment of DEGs shared between *fc2* vs wt and *fc2 cry2* vs wt in constant (24h) light conditions

Table S7. All differentially expressed genes identified between *fc2 cry2* and wt under cycling (6h light / 18h dark) light conditions

Table S8. Gene Ontology term enrichment of DEGs shared between *fc2* vs. wt and *fc2 cry2* vs wt in cycling (6h light / 8h dark) light conditions

Table S9. Gene Ontology term enrichment of DEGs specific to *fc2* vs. wt and not shared with *fc2 cry2* vs wt in cycling (6h light / 8h dark) light conditions

Table S10. Gene Ontology term enrichment of DEGs specific to *fc2 cry2* vs. wt and not shared with *fc2* vs. wt in constant (24h) light conditions

Table S11. Gene Ontology term enrichment of DEGs shared between *fc2 cry2* vs. wt in constant (24h) light conditions and *fc2* vs. wt in cycling (6h light / 18h dark) light conditions, but not *fc2* vs. wt in 24h light conditions

Table S12. All differentially expressed genes identified between *cry2* and wt under constant (24h) light conditions

Table S13. Gene Ontology term enrichment of DEGs identified in *cry2* vs. wt in constant light conditions

Table S14. All differentially expressed genes identified between *fc2 cry2* and *fc2* under cycling light (6h light / 18h dark) conditions

Table S15. Gene Ontology term enrichment of DEGs identified between *fc2 cry2* and *fc2* in cycling light (6h light / 18h dark) light conditions

Table S16. Opposing expression responses between *fc2* and *fc2 cry2* (relative to wt) in cycling light (6h light / 18h dark) conditions

Table S17. Gene Ontology term enrichment of DEGs identified with opposing expression responses between *fc2* and *fc2 cry2* (relative to wt) in cycling light (6h light / 18h dark) conditions

Table S18. Comparison of DEGs identified in this study with CRY2-regulated / blue-light independent DEGs identified in Zeng et al. 2025, *Cell*

Table S19. Comparison of DEGs identified in this study with CRY1-regulated / singlet oxygen-dependent DEGs identified in *flu*

Table S20. Mutant lines used for this study

Table S21. Primers used for RT-qPCR, genotyping, and sequencing

Table S22. Vectors used in this study

