## Supplemental materials (Figures S1-13 and Tables S20-22) for "Cryptochrome 2 is necessary for promoting a blue light-independent chloroplast signal to control cellular degradation"

### Supplementary Figures

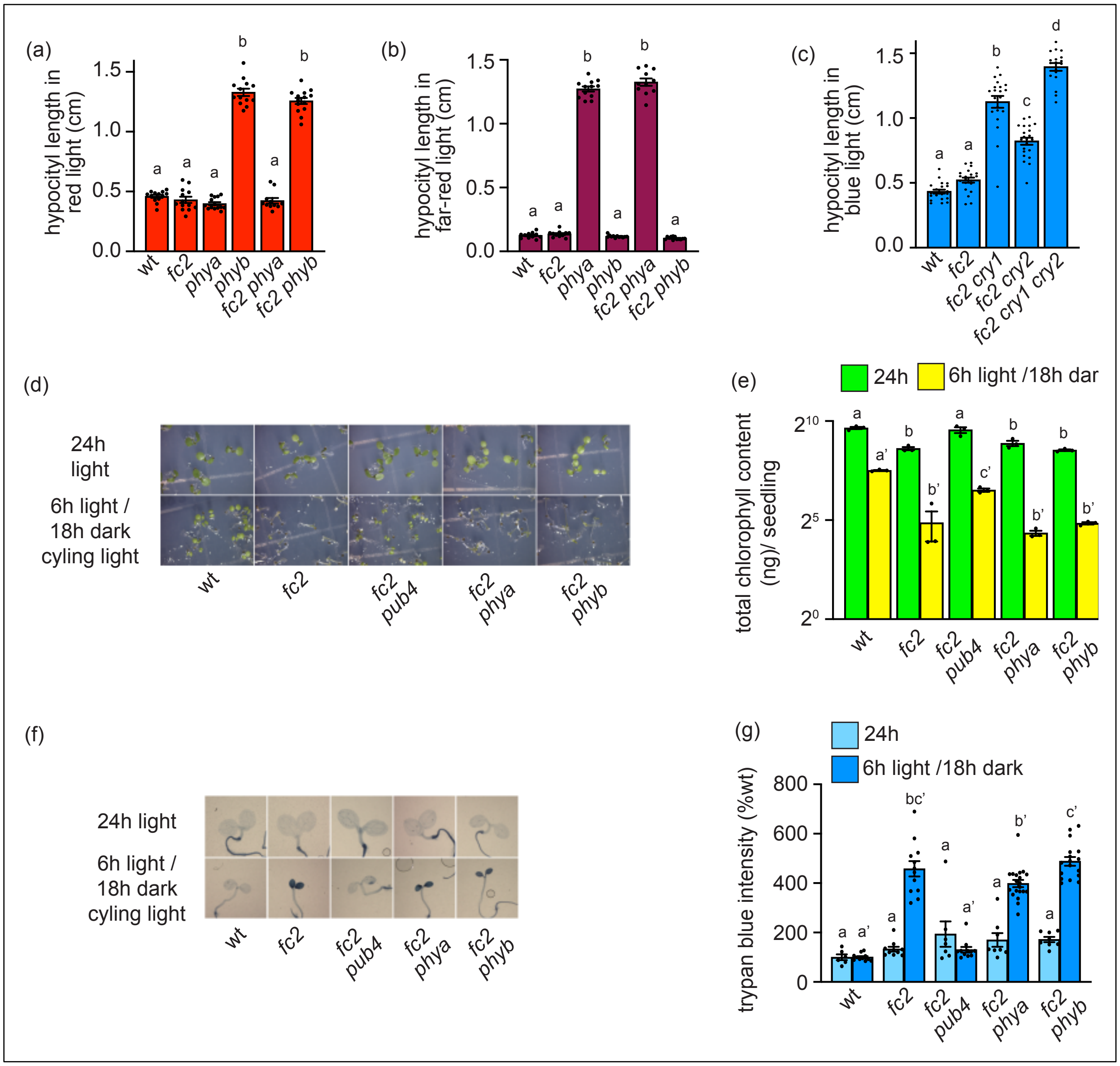


<u>Figure S1. phyA and phyB are not required for singlet oxygen induced cell death in *fc2* seedlings.</u> The effects of phytochrome mutations in the *fc2* genetic background was assessed in seedlings. **A)** Quantification of hypocotyl length of five-day old seedlings grown under constant red light conditions (8 µmol photons $m^{-2}$ $sec^{-1}$) (+/- SE, n $\geq$ 11). **B)** Quantification of hypocotyl length of five-day old seedlings grown under constant far-red light conditions (12 µmol photons $m^{-2}$ $sec^{-1}$) (+/- SE, n $\geq$ 11). **C)** Quantification of hypocotyl length of five-day old seedlings grown under constant blue light conditions (2 µmol photons $m^{-2}$ $sec^{-1}$ (+/- SE, n $\geq$ 18). **D)** Shown are six-day-old seedlings grown under constant white light (24h) or diurnal cycling white light (6h light / 18h dark) conditions. **E)** Mean levels of total chlorophyll (per seedling) of seven-day-old seedlings grown in 24h light (+/- SE, n = 3 replicates). **F)** Shown are representative images of seedlings from panel D stained with trypan blue. The dark blue color is indicative of cell death. **G)** Shown are mean intensities of trypan blue from panel F (+/- SE, n ≥ 6 seedlings). Statistical analyses in all bar graphs were performed using a one-way ANOVA followed by a Tukey HSD test. Different

letters above bars indicate significant differences ($p \leq 0.05$). Closed circles represent individual data points.

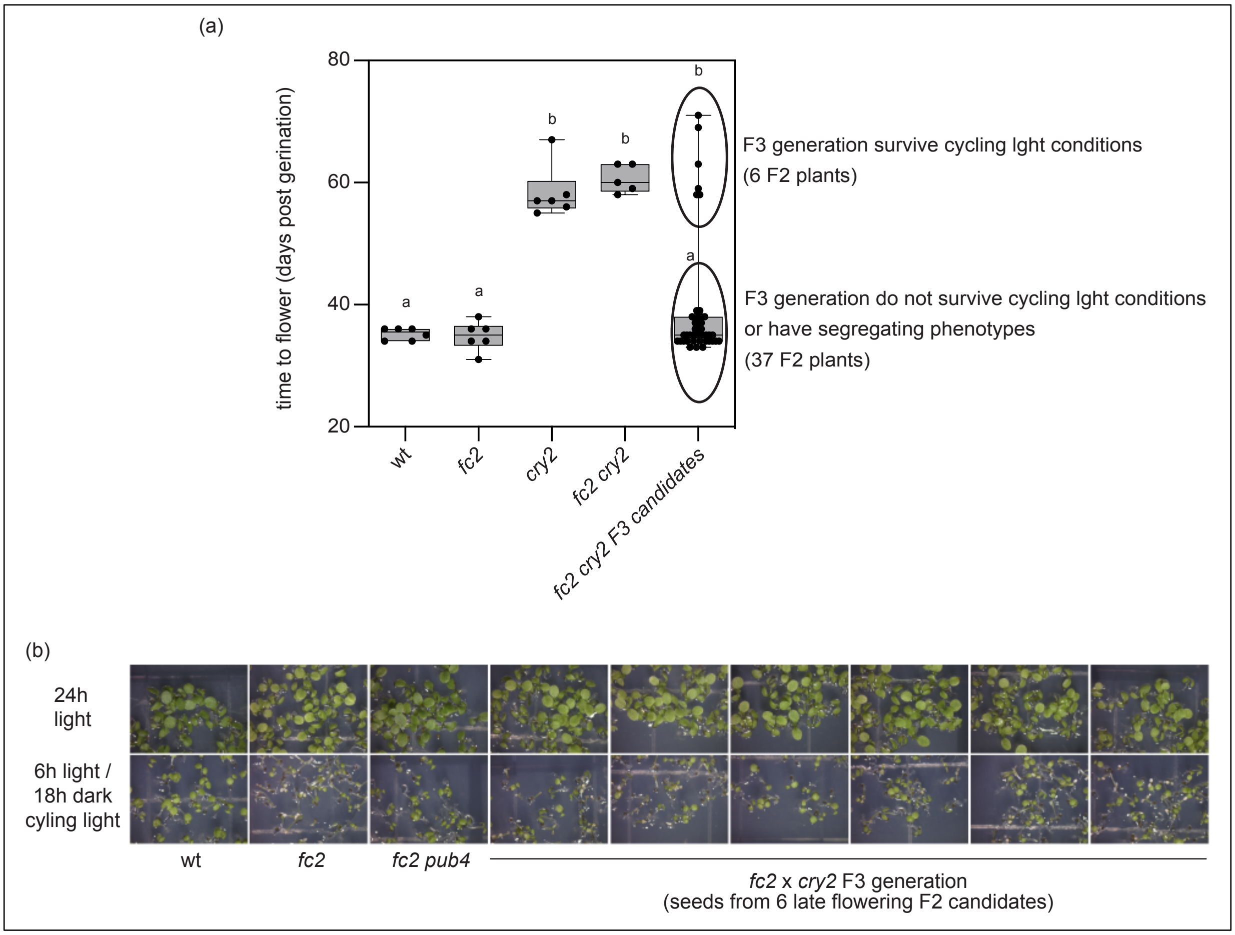


Figure S2. Late flowering and suppression of cell death phenotypes are linked in *fc2 cry2*.
A genetic linkage experiment was performed against the late flowering and suppression of cell death phenotypes in *fc2 cry2* mutant. **A)** Box and whisker plot (min to max) showing flowering time (days post germination) of the indicated genotypes as well as 43 *fc2*/*fc2* F2 plants from a *fc2*(♂) x *cry2*(♀) cross. Seeds were collected from all 43 plants and their phenotypes under cycling light conditions was assessed in the F3 generation. Of the 6 lines that were late to flower in the F2 generation, all suppressed cell death under cycling light conditions in the F3 generation. All lines were also genotyped to be homozygous for the *cry2-1* mutation. The 37 lines with wt-like flowering time in the F2 generation did not lead to suppression of cell death in the F3 generation, or had a segregating fts phenotype (1:3). These lines were genotyped and were *CRY2*/*CRY2* or *cry2-1*/*CRY2*. **A)** Shown are six-day-old seedlings grown under constant white light (24h) or diurnal cycling white light (6h white light / 18h dark) conditions.

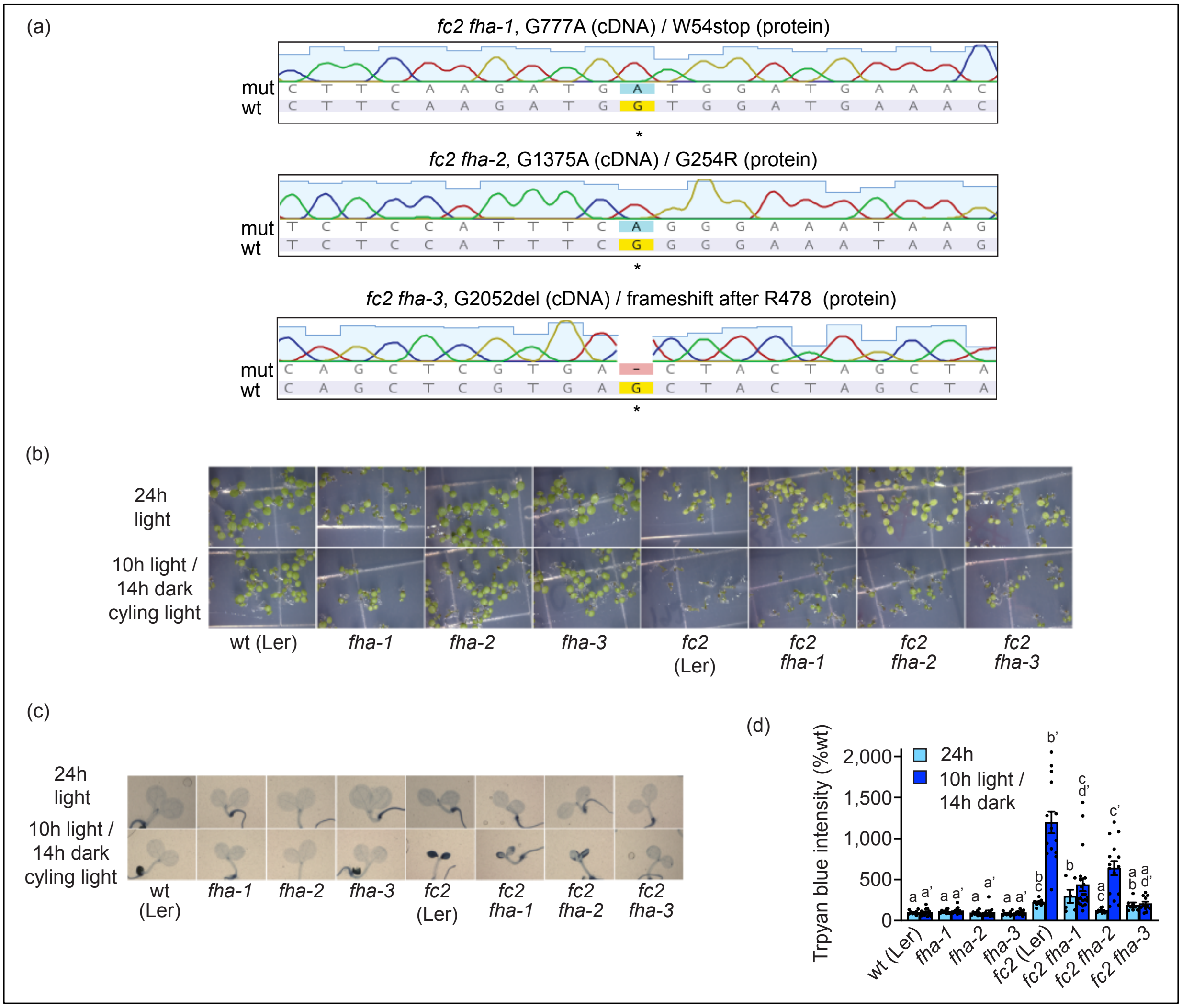


<u>Figure S3. *cry2* mutant alleles suppress cell death in the Landsberg erecta *fc2* mutant.</u>
*cry2* alleles in the Landsberg erecta (Ler) genetic background (*fha-1*, *fha-2*, and *fha-3*) were tested for their ability to suppress cell death in the Ler *fc2* mutant. **A)** *CRY2* sequences obtained from *fc2 fha-1*, *fc2 fha-2*, and *fc2 fha-3*. Arrow signifies the known mutation. Additional details are in **Table S1**. **B)** Shown are six-day-old seedlings grown under constant (24h) or diurnal light cycling (10h white light / 14h dark) conditions. **C)** Shown are representative images of the seedlings in panel B stained with trypan blue. The dark blue color is indicative of cell death. **D)** Shown are mean intensities of trypan blue (+/- SE, n ≥6 seedlings) from panel C. Statistical analyses in panel D were performed using a one-way ANOVA followed by a Tukey HSD test. Letters indicate statistically significant differences between samples ($P \le 0.05$). Separate analyses were performed for the different light conditions, and the significance of the cycling light is denoted by letters with a prime symbol ('). Closed circles represent individual data points.

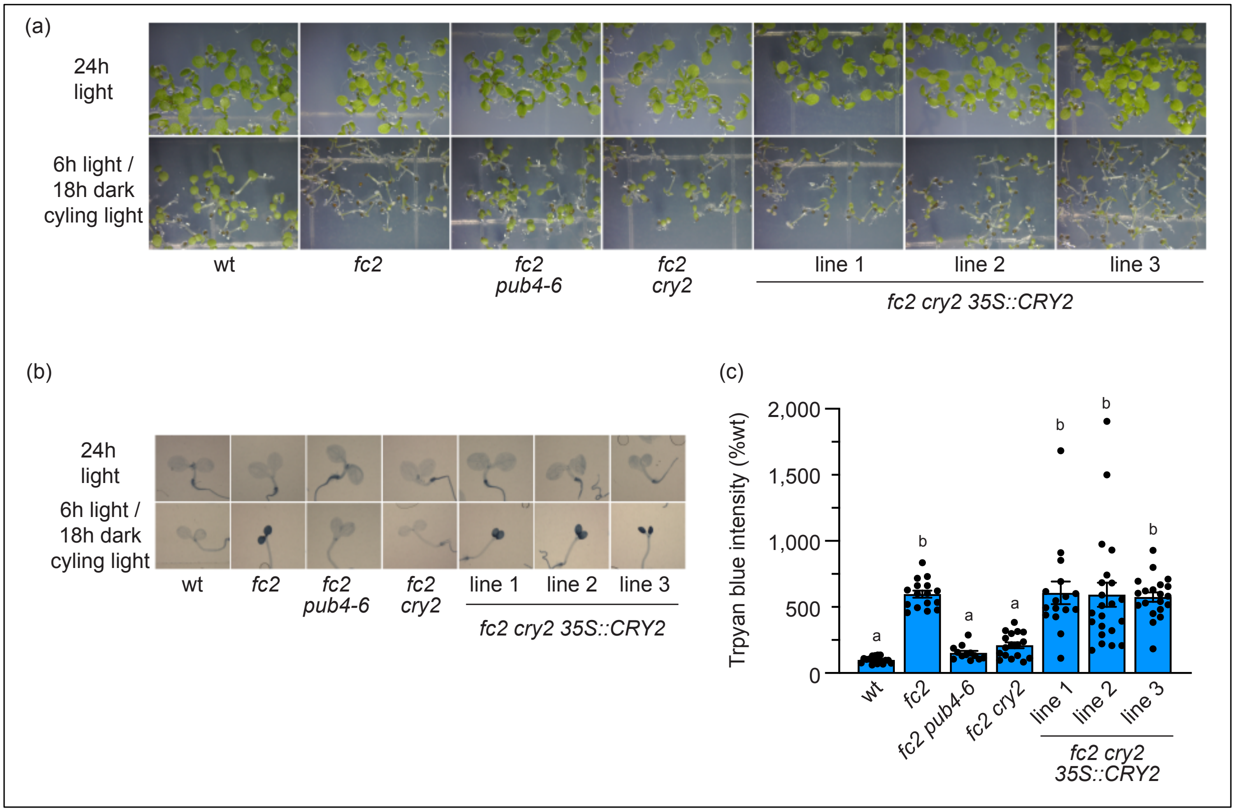


Figure S4. Complementation of the *cry2* mutation.
*fc2 cry2* mutants were complemented with a wt type copy of *CRY2* (*35S*::*CRY2*) to test causality of the cell death suppression phenotype in *fc2 cry2*. **A)** Shown are six-day-old seedlings grown under constant (24h) or diurnal cycling (6h white light / 18h dark) light conditions. **B)** Shown are representative images of the seedlings in panel A stained with trypan blue. The dark blue color is indicative of cell death. **C)** Shown are mean intensities of trypan blue intensities (+/- SE, n ≥ 10 seedlings) from panel B. Statistical analyses in **C** were performed using a one-way ANOVA followed by a Tukey HSD test. Letters indicate statistically significant differences between samples ($P \leq 0.05$). Closed circles represent individual data points.

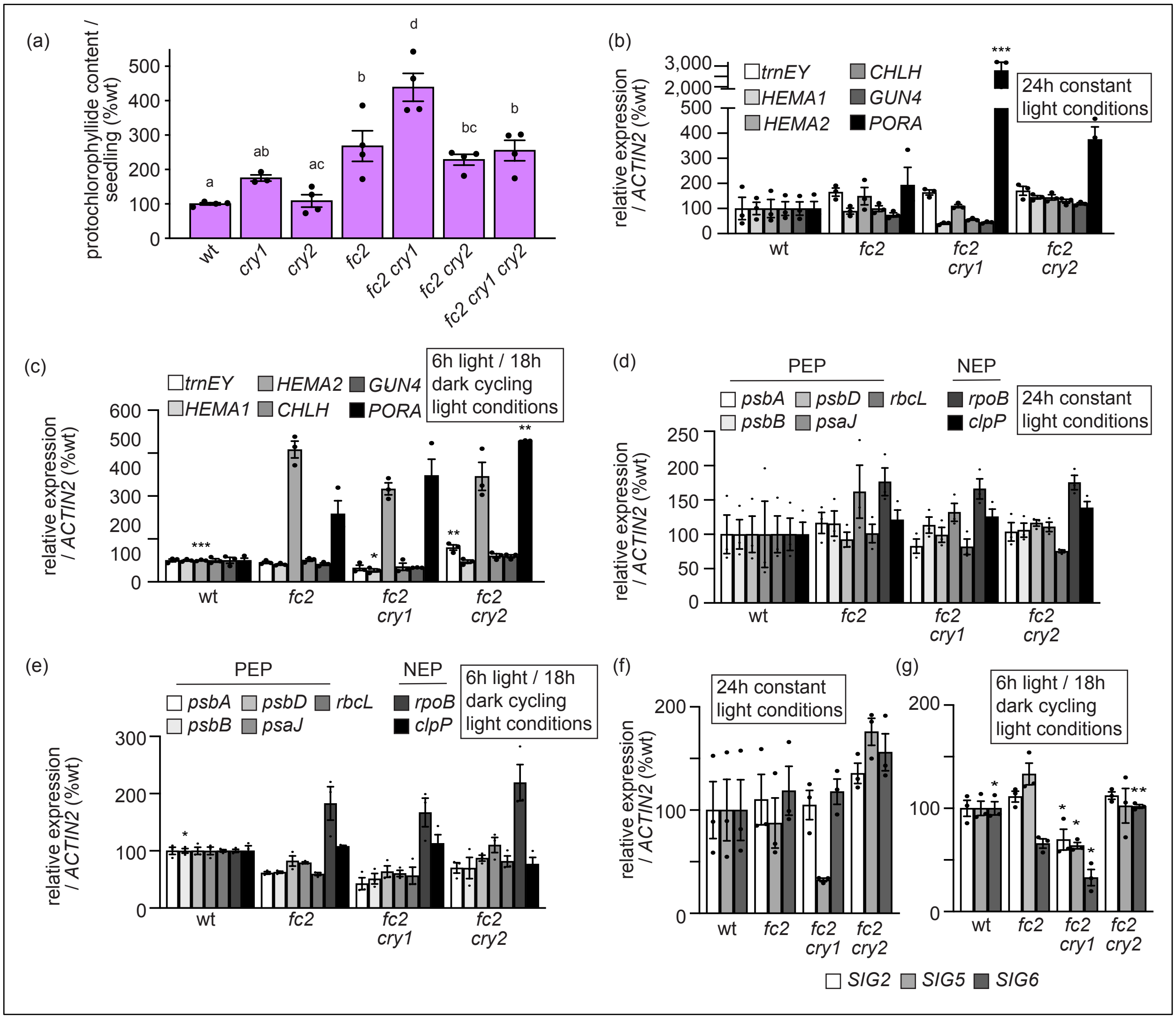


Figure S5. Testing the impact of *cry2* on chloroplast development.

The impact of the *cry1* and *cry2* mutations on chloroplast development in *fc2* seedlings was determined. **A)** Mean levels of protochlorophyllide in six-day-old dark grown (etiolated) seedlings (+/- SE, n = 4 groups of 10 seedlings). RT-qPCR analysis of steady-state transcripts of **B)** and **C)** genes involved in tetrapyrrole metabolism, **D)** and **E)** plastid-encoded genes, and **F)** and **G)** *SIGMA FACTOR* (*SIG*) genes. Panels **B**, **D**, and **F** show expression from four-day-old seedlings grown under 24h constant light conditions. Panels **C**, **E**, and **G** show expression from four-day-old seedlings grown under diurnal light cycling conditions (6h light / 18h dark). Shown are mean expression values (+/- SE, n = 3 biological replicates). Statistical analyses in performed using a one-way ANOVA followed by a Tukey HSD test. In panel A, letters indicate statistically significant differences between samples ($P \leq 0.05$). In panels B-F, statistical significance in respect to *fc2* is indicated as follows: * = p-value ≤ 0.05, ** = p-value ≤ 0.01, *** = p-value ≤ 0.001. Closed circles represent individual data points.

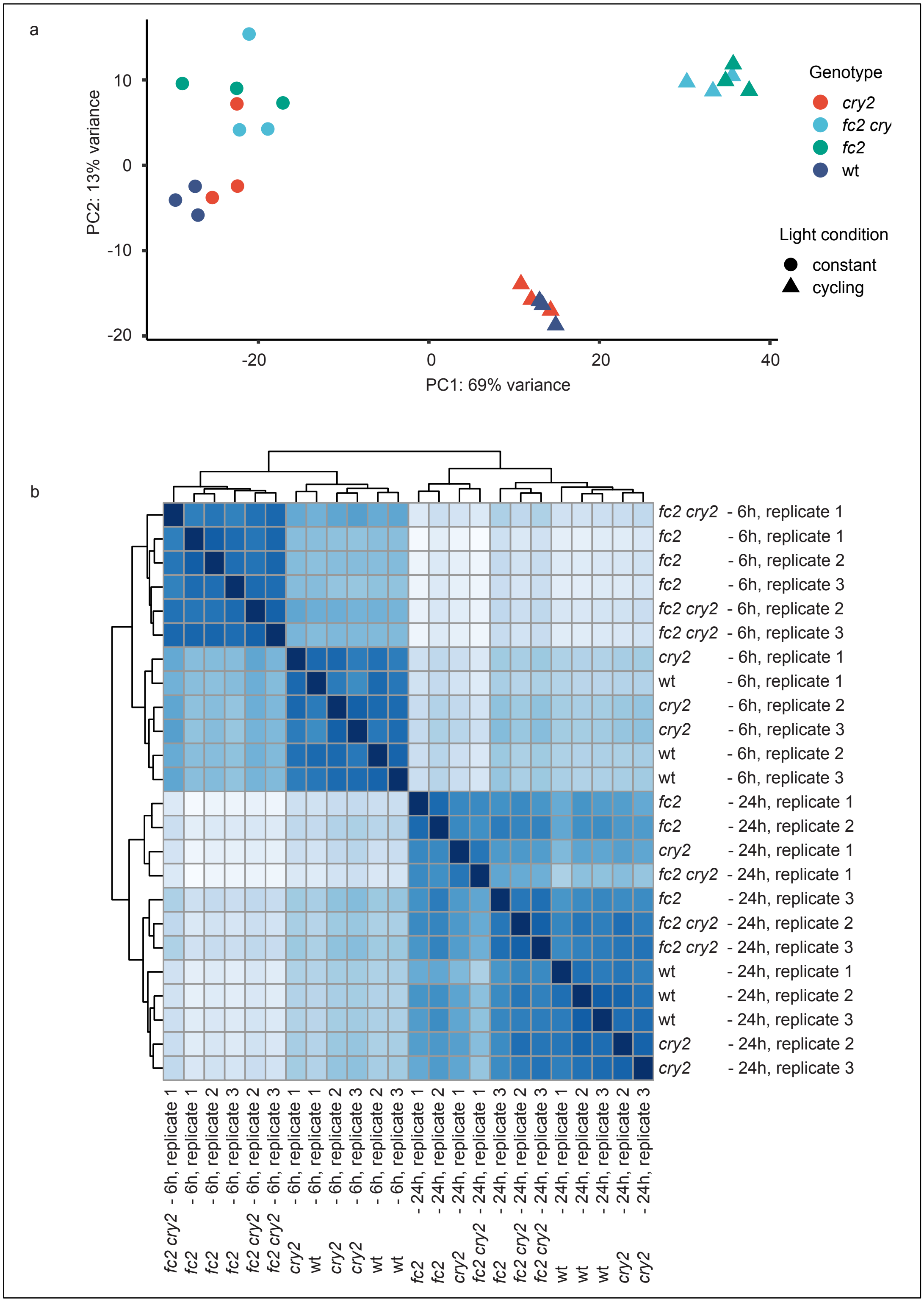


Figure S6. Visualization of variance among RNA-seq replicates used in the study.
Shown are two analyses to visualize grouping and similarity among the 24 RNA-seq replicates used in this study to assess data quality. Four genotypes (wt, *fc2*, *cry2*, and *fc2 cry2*) were grown

for four days in two conditions (constant light conditions (24h) or 6h light / 18h dark diurnal cycling light conditions (cycling)) for a total of eight groups with three replicates each. **A)** Principal component analysis (PCA) scatter plot showing separation of the RNA-seq samples relative to each other. The genotypes and conditions are color and shape coded, respectively, according to the key on the right. PCA was performed using the DESeq2 R package on Salmon imported quantification data using default parameters. Variance explained for each PC (PC1: 69% variance, PC2: 13% variance) is reported in brackets with the axis label. **B)** Shown is a sample-to-sample distance heatmap for the same 24 replicates. Sample distances were calculated using the "dist" function in R on the DESeq2 object. The heatmap was then plotted using Pheatmap in R to represent relative similarity among the individual RNA-seq samples. Shades of blue indicate distance between samples with darker shades indicating smaller distance and lighter shades indicating greater distance.

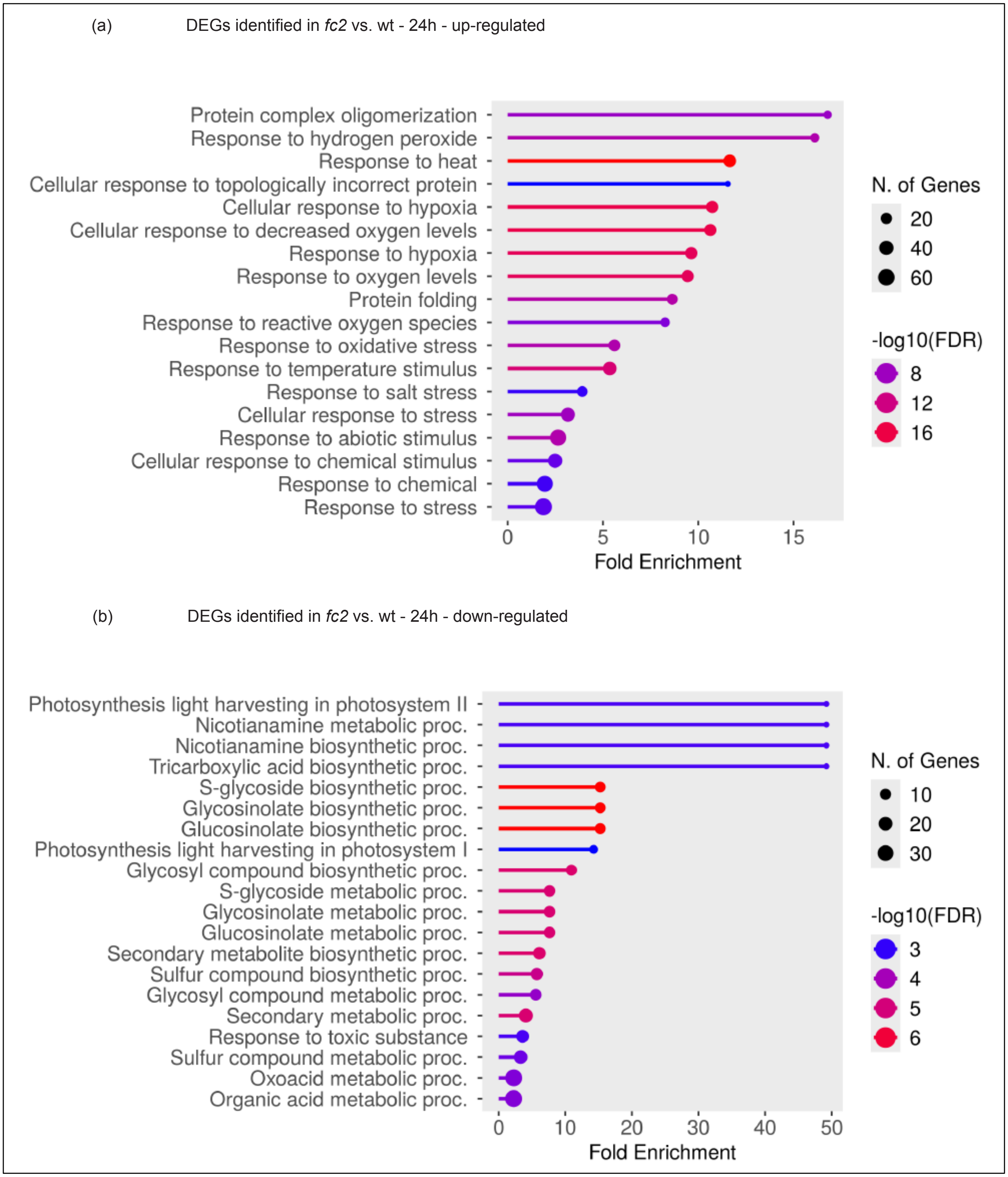


<u>Figure S7. Gene ontology analyses of differentially expressed genes in *fc2* vs. wt under constant light conditions.</u>

Shown are gene ontology (GO) term enrichment analyses of the identified differentially expressed genes (DEGs) in this study. **A)** GO-term enrichment from the 228 up-regulated DEGs from *fc2* vs. wt under constant (24h) light conditions. **B)** GO-term enrichment from the 313 down-regulated

DEGs from the same comparison. GO-term enrichment analyses were performed using ShinyGO 0.82 with an FDR cutoff of p ≤ 0.05. x-axes indicate fold-enrichment. The ball size indicates number of genes. The line colors represent FDR values.

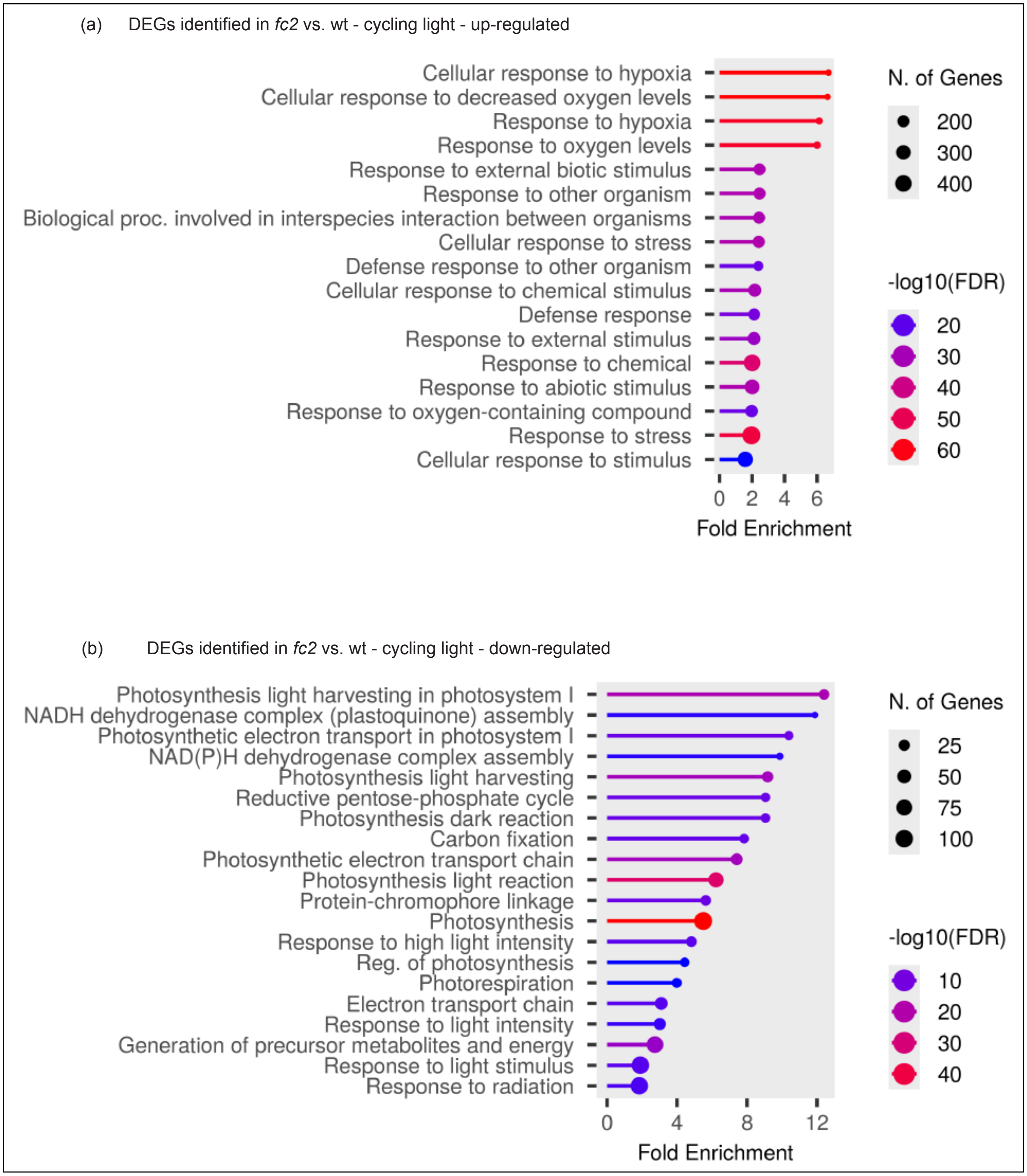


<u>Figure S8. Gene ontology analyses of differentially expressed genes in *fc2* vs. wt under diurnal cycling light conditions.</u>
Shown are gene ontology (GO) term enrichment analyses of the identified differentially expressed genes (DEGs) in this study. **A)** GO-term enrichment from the 1563 up-regulated DEGs from *fc2* vs. wt under diurnal cycling light (6h light / 18h dark) conditions. **B)** GO-term enrichment from the 1524 down-regulated DEGs from the same comparison. GO-term enrichment analyses were

performed using ShinyGO 0.82 with an FDR cutoff of $p \leq 0.05$. x-axes indicate fold-enrichment. The ball size indicates number of genes. The line colors represent FDR values.

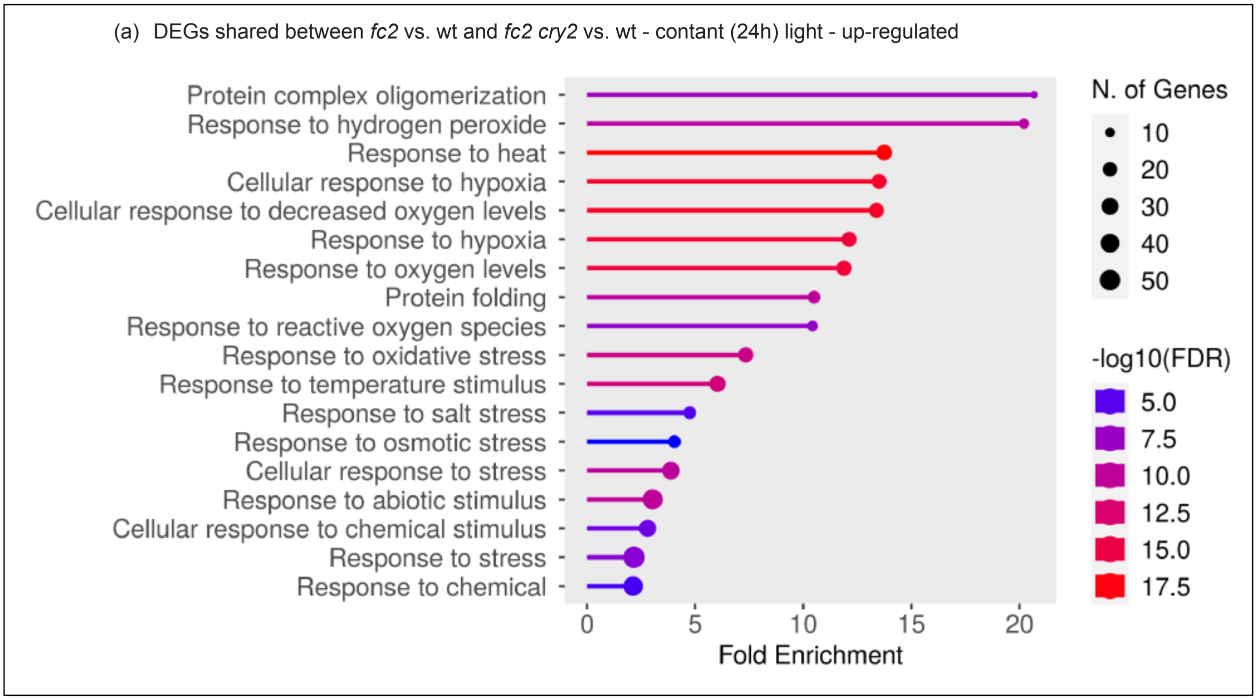


Figure S9. Gene ontology analyses of differentially expressed genes shared between *fc2* vs. wt and *fc2 cry2* vs. wt under constant light conditions.

Shown are gene ontology (GO) term enrichment analyses of the identified differentially expressed genes (DEGs) in this study. **A)** GO-term enrichment from the 164 up-regulated DEGs shared between comparisons of *fc2* vs. wt and *fc2 cry2* vs. wt under constant light (24h) conditions. GO-term enrichment analyses were performed using ShinyGO 0.82 with an FDR cutoff of $p \leq 0.05$. x-axes indicate fold-enrichment. The ball size indicates number of genes. The line colors represent FDR values.

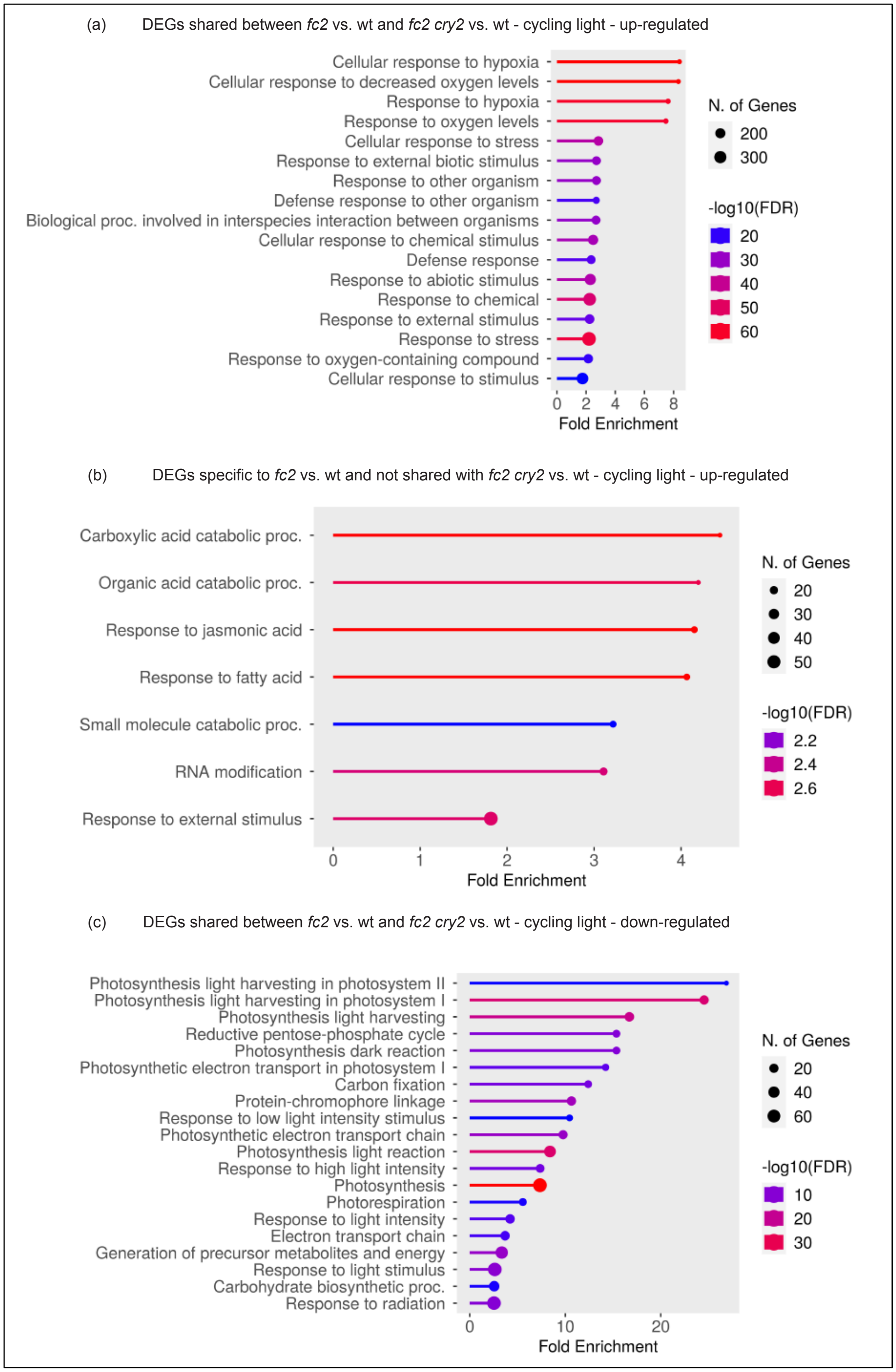


<u>Figure S10. Gene ontology analyses of differentially expressed genes identified in *fc2* vs. wt under cycling light conditions.</u>

Shown are gene ontology (GO) term enrichment analyses of the identified differentially expressed genes (DEGs) in this study. **A)** GO-term enrichment from the 1127 up-regulated DEGs shared between comparisons of *fc2* vs. wt and *fc2 cry2* vs. wt under cycling light (6h light / 18h dark) conditions. **B)** GO-term enrichment from the 436 up-regulated DEGs identified in *fc2* vs. wt in cycling light conditions that are not shared with *fc2 cry2* vs. wt in cycling light conditions. **C)** GO-term enrichment from the 773 down-regulated DEGs shared between comparisons of *fc2* vs. wt and *fc2 cry2* vs. wt under cycling light (6h light / 18h dark) conditions. GO-term enrichment analyses were performed using ShinyGO 0.82 with an FDR cutoff of $p \leq 0.05$. x-axes indicate fold-enrichment. The ball size indicates number of genes. The line colors represent FDR values.

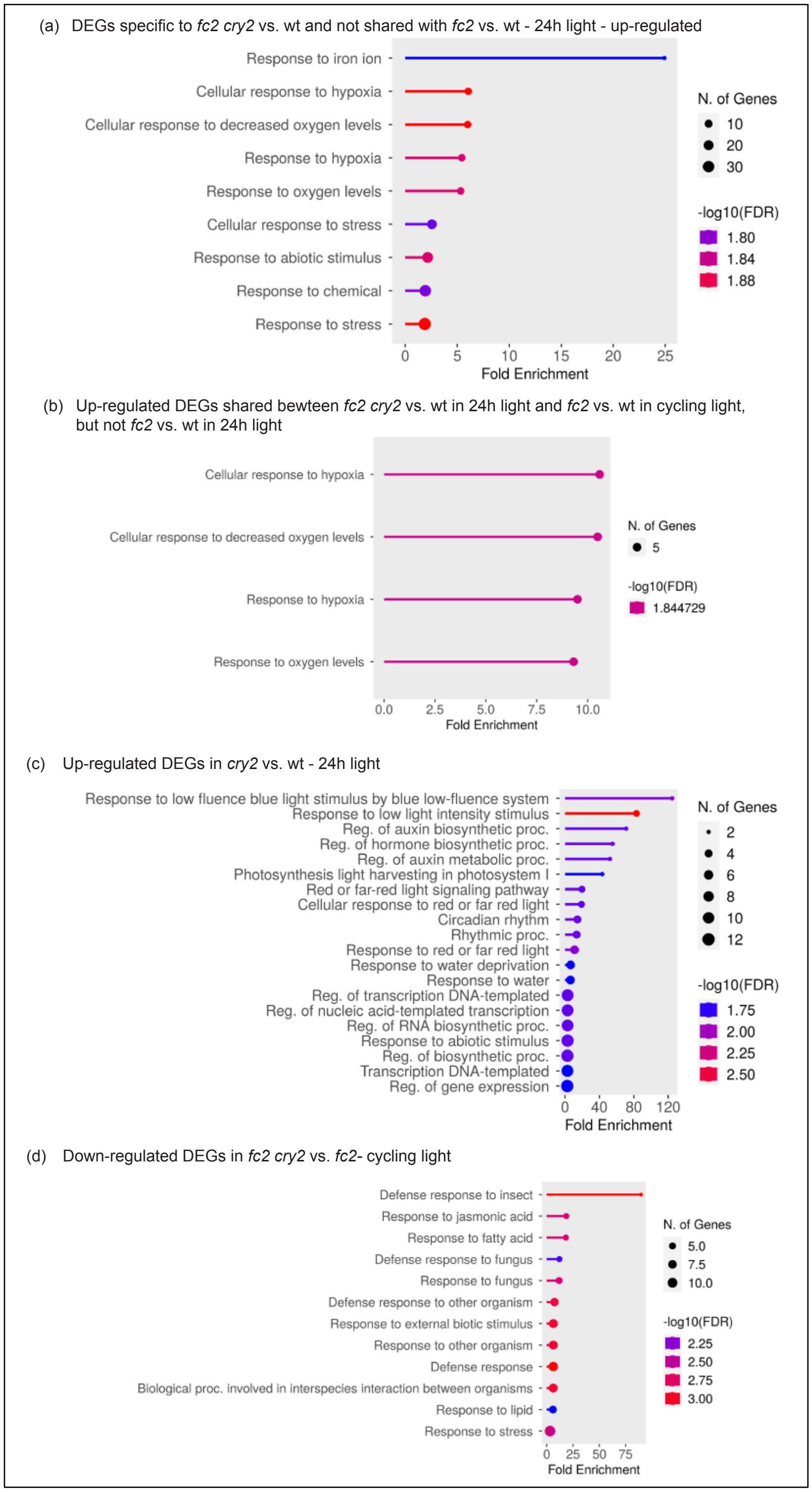


<u>Figure S11. Gene ontology analyses of differentially expressed genes identified in *cry2* and *fc2 cry2* mutants.</u>

Shown are gene ontology (GO) term enrichment analyses of the identified differentially expressed genes (DEGs) in this study. **A)** GO-term enrichment from the 132 up-regulated DEGs identified in *fc2 cry2* vs. wt under constant light (24h) conditions that are not shared with *fc2* vs. wt in constant light conditions. **B)** GO-term enrichment from the 48 up-regulated DEGs identified in *cry2 fc2* vs. wt in constant light conditions, shared with *fc2* vs. wt in cycling light (6h light / 18h dark) conditions, but not shared with *fc2* vs. wt in constant light conditions. **C)** GO-term enrichment from the 43 up-regulated DEGs identified in *cry2* vs. wt under constant light conditions. **D)** GO-term enrichment from the 27 down-regulated DEGs identified between in *fc2 cry2* vs. *fc2* under cycling light conditions. GO-term enrichment analyses were performed using ShinyGO 0.82 with an FDR cutoff of $p \leq 0.05$. x-axes indicate fold-enrichment. The ball size indicates number of genes. The line colors represent FDR values.

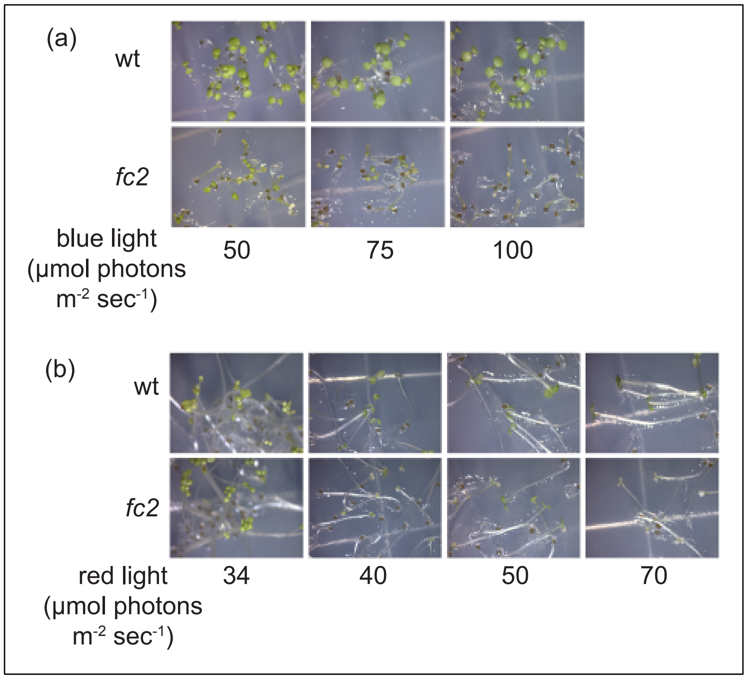


Figure S12. Testing level of red and blue light necessary to induce cell death in *fc2* mutant seedlings.

Various levels of monochromatic blue (bc) and red light (rc) were tested to determine the amount necessary to induce cell death in fc2 seedlings during cycling light (6h light / 18h dark) conditions. **A)** Shown are six-day-old seedlings grown under diurnal cycling rc (6h rc / 18h dark) using the indicated μmol photons $m^{-2}$ $sec^{-1}$. **B)** Shown are six-day-old seedlings grown under diurnal cycling bc (6h bc / 18h dark) using the indicated μmol photons $m^{-2}$ $sec^{-1}$.

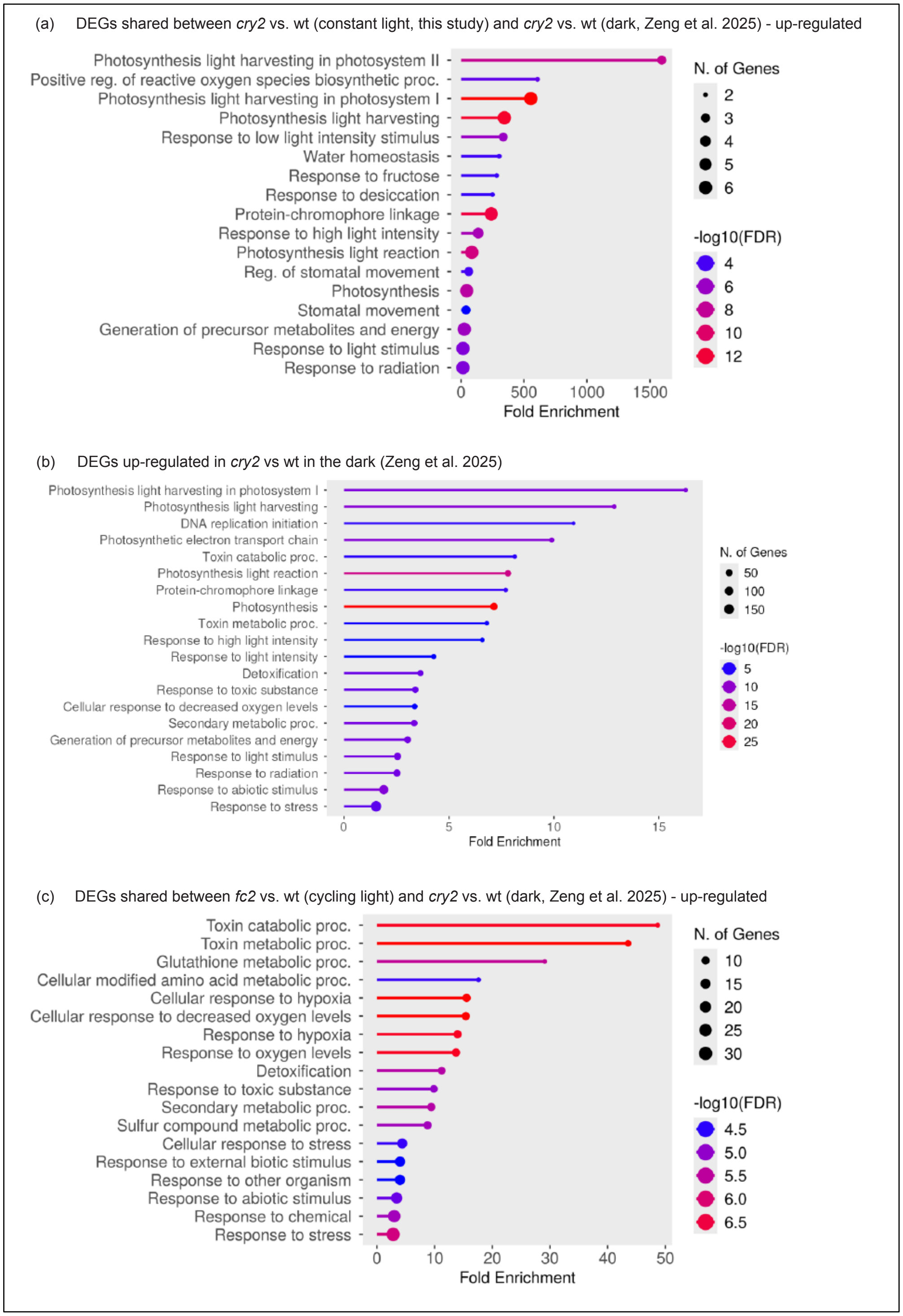


<u>Figure S13. Gene ontology analyses of blue light-independent differentially expressed genes identified in *cry2* and *fc2* mutants</u>

Shown are gene ontology (GO) term enrichment analyses of the identified differentially expressed genes (DEGs) in this study and in [1]. **A)** GO-term enrichment from the 13 up-regulated DEGs shared between *cry2* vs. wt under constant light (24h, this study) and *cry2* vs wt in the dark [1]. **B)** GO-term enrichment from the 834 up-regulated DEGs identified in *cry2* vs. wt in the dark [1]. **C)** GO-term enrichment from the 76 up-regulated DEGs shared between in *fc2* vs. wt under cycling light conditions and *cry2* vs. wt in the dark [1]. GO-term enrichment analyses were performed using ShinyGO 0.82 with an FDR cutoff of $p \leq 0.05$. x-axes indicate fold-enrichment. The ball size indicates number of genes. The line colors represent FDR values.

### **Supplementary Tables**

Table S20. Mutant plant lines used for this study.

| Mutant | Gene Name | Gene # | Mutation | Notes | Ref |
|---|---|---|---|---|---|
| *fc2-1* | *PLASTID FERROCHELATASE 2* | *At2g30390* | GABI_ 766H08 T-DNA in 5'UTR | Sulfadiazine Resistant | [2] |
| *cry1-304* | *CRYPTOCHROME 1* | *At4g08920* | Deletion; Loss of Function | Neutron Mutagenesis | [3] |
| *cry2-1* | *CRYPTOCHROME 2* | *At1g04400* | Deletion; Loss of Function | Neutron Mutagenesis | [4] |
| *phyA-211* | *PHYTOCHROME A* | *At1g09570* | Unknown; Loss of Function | Gamma Radiation | [5] |
| *phyB-9* | *PHYTOCHROME B* | *At2g18790* | W397STOP; Loss of Function | Ethyl methanesulfonate | [6] |
| *fha-2* | *CRYPTOCHROME 2* | *At1g04400* | G254R | Ethyl methanesulfonate | [7] |
| *fha-1* | *CRYPTOCHROME 2* | *At1g04400* | W54STOP | Ethyl methanesulfonate | [7] |
| *fha-3* | *CRYPTOCHROME 2* | *At1g04400* | Deletion | X-ray | [7] |
| *pub4-6* | *PLANT U-BOX 4* | *At2g23140* | G255R | EMS | [8] |

Table S21. Primers used for RT-qPCR, genotyping, and sequencing

| Gene | Primer orientation / name | Sequence |
|---|---|---|
| **RT-qPCR primer pairs** | | |
| *ACTIN2 / At3g18780* | For. / JP199 | GCACTTGCACCAAGCAGCAT |
| | Rev. / JP200 | CCTTTCAGGTGGTGCAACGAC |
| *LHCB1.2 / CAB3 / At1g29910* | For. / JP197 | GGACTTGCTTTACCCCGGTG |
| | Rev. / JP198 | TCGGTAGCAAGACCCAATGG |
| *LHCB 2.2 / At2g05070* | Rev. / WLO1529 | GCTTTGTAAACTCGTGATTGTG |
| | For. / WLO1530 | TGCCAAATTCACATCAAACG |
| *RBCS 2B / At5g38420* | Rev. / WLO1448 | GCTTCACCGAAGCTTAATCC |
| | For. / WLO1449 | CCACATAGAAATGGGTTCCAG |
| *CA1 / At3g01500* | For. / JP209 | TGTGTCCATCACACGTTCTGG |
| | Rev. / JP210 | GGACCACGAAGGCATCTCCT |
| *psaJ / AtCg00630* | Rev. / WLO1539 | GTACTCTATGGTTCGGTTCGT |
| | For. / WLO1540 | AGGGAAATGTTAATGCATCTGG |
| *psbA / AtCg00020* | For. / WLO1535 | CGTCTTTACATTGGATGGTTTGG |
| | Rev. / WLO1536 | CAGAAGTTGCGGTCAATAAGG |
| *psbB / AtCg00680* | For. / WLO1537 | TCGTGCGACTTTGAAATCTGA |
| | Rev. / WLO1538 | CAACCTCTTGGGCTGCTACG |
| *psbD / AtCg00270* | For. / WLO1969 | TGCAATCGCATTCTCTGGTC |
| | Rev. / WLO1970 | ACTAGGCGCAAAGAACCAAC |
| *rbcL / AtCg00490* | For. / WLO1541 | TTACAAAGGACGATGCTACCACAT |
| | Rev. / WLO1542 | TGAGTTTCTTCTCCTGGAACGG |
| *clpP / AtCg00670* | For. / WLO1545 | TGGGTTGACATATACAACCGACTTT |
| | Rev. / WLO1546 | GCCTAAAAAAAATAATCTTTCTCGATAA A |
| *rpoB / AtCg00190* | For. / WLO1547 | ATACGAGATATCCATCCTAGTCAC |
| | Rev. / WLO1548 | GTCCAACATTGATTCCTTCAGAC |
| *trnEY / AtCg00250-240* | For. / WLO1549 | TCTAGTGGTTCAGGACATCTC |
| | Rev. / WLO1550 | TTGCCAACGAATTTACAGTC |
| *HEMA1 / At1g58290* | For. / JP254 | GCTTCCGCAGTCTTCAAACG |
| | Rev. / JP255 | CCAGCGCCAATTACACACATC |
| *HEMA2 / At1g04490* | For. / JP211 | AGCTCCTGCACGGTCCAAT |
| | Rev. / JP212 | TGCTATCGTTCCCATCGCAT |
| *CHLH / At5g13630* | For. / WLO1692 | TGCAAGCTTTGGAAGGCAAG |
| | Rev. / WLO1693 | ACTTGCCATTGCTGCTGTTG |
| *GUN4 / At3g59400* | For. / WLO1628 | ATTCGGATACAGCGTGCAAC |
| | Rev. / WLO1629 | ATTCGTCAGGAAACGCTCTG |
| *PORA / At5g54190* | For. / WLO1626 | AGCAACGGCAAAGGCATTAG |
| | Rev. / WLO1627 | ACGCCAAGTCCAAATGCATC |
| *SIB1 / At3g56710* | For. / JP589 | CAACCGGAGCCCATCTATT |
| | Rev. / JP590 | GGAGAAAGGTTGTGGTCGTC |
| *HSP26.5 / At1g52560* | For. / JP585 | CGAGCTTATCGTTGCCTGAT |
| | Rev. / JP586 | CTCCGCCTTAATGTCCTCAA |
| *BAP1 / At3g61190* | For. / JP338 | ATTGATGGATACGGTGGCCG |

| | | |
|---|---|---|
| | Rev. / JP339 | CAGACCCCAAACCGGAACTC |
| *Atpase / At3g28580* | For. / JP336 | GAAGATCGGAAAAGCGTGGAA |
| | Rev. / JP337 | CCGGGTGGTCCAAACAAAAG |
| *ZAT12 / At5g59820* | For. / JP344 | GCGTTGGTTACACGCGCTT |
| | Rev. / JP345 | CTTCAACGTAGTCACCGTGGG |
| *CYC8 / At4g37370* | For. / JP1130 | AATGGGCATTGTCGAACGTG |
| | Rev. / JP1131 | TCGCCTTGTTCAATACATCCG |
| *CRY1 / At4g08920* | For. / WLO1487 | ACTGGTTGGTTGCATGATCG |
| | Rev. / WLO1488 | ATTGCCAACCAAGAGCATCG |
| *CRY2 / At1g04400* | For. / WLO1489 | AATAACTGCAGCGGCTGAAG |
| | Rev. / WLO1490 | ACAACGCATTGCTCGGTTTC |
| **Genotyping primers (fragment size)** | | |
| *fc2-1* (GabiKat_766H08) (~600bp) | For. / JP283 | GAGCAACGCCAAACATAGAAG |
| | Rev. / JP285 | TCAAAGGCAATGAATGTTTCC |
| *wt FC2* (~1kb) | For. / JP283 | GAGCAACGCCAAACATAGAAG |
| | Rev. / JP284 | TCAAAGGCAATGAATGTTTCC |
| *cry1-304* (~1.25kb) | For. / WLO1451 | ATGTCTGGTTCTGTATCTGGTTGTGGTTC |
| | Rev. / WLO1452 | ATAGTTCTCATCCACAGCCCAAG |
| *cry2-1* (~700bp) | For. / WLO1527 | CACCGTCTCATAATCTATAC |
| | Rev. / WLO1528 | CCAGTTTATCCTTTAAGCCTCATATCG |
| Vector pWLP396 (~3kb) | For. / WLO2304 | GACTATAGTTTGGTTTAGAAGAGACCT |
| | Rev. / WLO2305 | GGTACCGTCGACTGCAGAAT |
| Vector FGFP (~650bp) | For. / WLO2316 | TCGGCATGGACGAGCTGTA |
| | Rev. / WLO2317 | AAGCCGACTGCACTATAGCA |
| Vectors FGFP-CRY2 FGFP-CRY2$^{D387A}$ FGFP-CRY2$^{P532L}$ (~3kb) | For. / WLO2316 | TCGGCATGGACGAGCTGTA |
| | Rev. / WLO2317 | AAGCCGACTGCACTATAGCA |
| **Sequencing primers** | | |
| *CRY2* | For. / WLO2012 | AGATGGACAAAAAGACTATAGTTTGGT |
| | For. / WLO2013 | CCCACAACACGATTTCAGCG |
| | For. / WLO2014 | CTCCGTATCTCCATTTCGGGG |
| | For. / WLO2015 | TGGACAATCCCGCGTTACAA |
| | Rev. / WLO2018 | ACTTCACAGCAAAGCTTGAAACA |
| | Rev. / WLO2054 | ACAACGCATTGCTCGGTTTC |
| | Rev. / WLO2055 | CGCTGAAATCGTGTTGTGGG |
| **Cloning primers** | | |
| *CRY2* cDNA coding region with stop codon | For. / WLO1904 | GGGGACAAGTTTGTACAAAAAAGCAGGCTTGATGAAGATGGACAAAAAGACT |
| | Rev. / WLO1906 | GGGGACCACTTTGTACAAGAAAGCTGGGTCTCATTTGCAACCATTTTTTCCCA |

| **Miscellaneous primers** | | |
|---|---|---|
| To amplify *CRY2* in *fha* mutants for sequencing | For. / WLO2046 | ATGAAGATGGACAAAAAGACT |
| | Rev. / WLO2047 | TCATTTGCAACCATTTTTTCCCA |

Table 22. Vectors used in this study

| Plasmid | Insert | Notes | Ref. |
|---|---|---|---|
| pWLP391 | *35S*::*CRY2* | pDONR221 | This study |
| pWLP396 | *35S*::*CRY2* | pEARLEYGATE101 | This study |
| FGFP | *ACT2*::*GFP* | | [9] |
| FGFP-CRY2 | *ACT2*::*GFP-CRY2* | | [9] |
| FGFP-CRY2$^{D387A}$ | *ACT2*::*GFP-CRY2*(D387A) | | [9] |
| FGFP-CRY2$^{P532L}$ | *ACT2*::*GFP-CRY2*(P532L) | | [9] |
